# Multi-omic Longitudinal Analysis of Canine Osteosarcoma Identifies Inter-Patient Heterogeneity and Immune Enrichment in Metastatic Lesions

**DOI:** 10.64898/2026.01.05.696411

**Authors:** Christopher Husted, Kerstin Seidel, Tanya T. Karagiannis, Cornelia Peterson, Kate Megquier, Diane Genereux, Jillian Richmond, Elinor Karlsson, David S. Shulman, Brian Crompton, Cheryl A. London, Heather L. Gardner

## Abstract

Osteosarcoma (OS) exhibits substantial genomic complexity and inter-patient heterogeneity, necessitating longitudinal, patient-matched analyses to understand acquired features of tumor evolution. However, most published OS data is limited to primary tumor samples, limiting insight into patient-specific resistance mechanisms. To address this, we characterized the genomic landscape of paired primary and metastatic tumor samples from dogs with spontaneous OS. Whole-genome and single-cell RNA sequencing reveal mutation and gene expression profiles that are predominantly organized by patient identity. Mutational burden and pathway alterations such as those involving PI3K, NOTCH, TP53, MAPK, RAS and epigenetic regulation differ between primary and metastatic samples. Variants present in tumor tissue are readily detectable in paired cfDNA samples, demonstrating the utility of this assay for identifying tumor-specific alterations associated with treatment resistance. Analysis of bulk RNA-seq data to estimate cell-type composition shows greater immune cell representation in metastases, underscoring the importance of immune signaling pathways in OS. These findings exemplify the presence of patient-specific alterations in genomic architecture over the course of tumor progression, linking CNV amplification, pathway reprogramming, and immune evasion in metastatic OS.

## Introduction

Osteosarcoma (OS) is the most common primary bone tumor in humans, and despite over four decades of research, treatment-resistant pulmonary metastatic disease remains the leading cause of death^1,2^. Notably, the five-year survival rate for localized disease has not improved beyond 60-70%, and this drops to 30% for those patients that present with metastasis at diagnosis^3–6^. To better understand the molecular and genomic features that drive OS biology, and thus improve therapeutic strategies, multiple sequencing datasets have been generated. However, given the challenges associated with routine collection of metastatic samples from adolescent patients, most of the available datasets are derived from primary tumors. Moreover, when primary and metastatic OS sequencing data are reported, they are often not patient-matched and may not include critical outcome linked information.

Understanding intra-patient alterations over the course of treatment and development of resistance in OS is critical for several reasons. OS is characterized by extreme genomic complexity, including widespread copy number alterations, chromothripsis (catastrophic chromosome shattering and rearrangement), and structural rearrangements ^7–13^, suggesting a high degree of genomic instability^14,15^. While large copy number alterations appear longitudinally stable in individual tumors ^14^, chromothripsis-driven tumor evolution may also contribute to the emergence of clonal drivers of metastasis^10^. Lastly, the microenvironment of the primary tumor (bone) is considerably different from that of the lung, the primary location for metastatic disease. As such, understanding how tumor cells evolve and interact in the context of the metastatic niche necessitates interrogation of changes in patient matched samples.

OS is the most common primary bone tumor in pet dogs, with lesions predominantly occurring in the appendicular skeleton. The therapeutic approach (surgical removal and chemotherapy) and development of metastasis in pet dogs parallel the disease course of OS in adolescent patients. Multiple groups have demonstrated high conservation of the OS tumor genome across pet dogs and humans, further underscoring the value of studying longitudinal tumor genomic and transcriptomic alterations in this species^16,17^. As the incidence of OS is 25-50 times higher than in humans and half of affected pet dogs develop lung metastasis within 6-9 months post tumor removal and chemotherapy, studies can be undertaken in a relatively compressed timeline. Critically, as most affected pet dogs with metastatic OS undergo planned euthanasia following clinical deterioration, patient matched primary and metastatic tumor samples with associated outcomes linked data can be more readily obtained.

Leveraging pet dog OS as a model for human OS, we hypothesized that (i) mutational profiles and cellular programs are more similar among samples from a given patient than between primary or metastatic tumors from different patients; (ii) relative to primary tumors, metastatic lesions exhibit increased genome-wide alteration burden and structural remodeling reflecting evolution of cell state programs; and (iii) cell-free (cf)DNA will provide a similar and complementary view of the genomic landscape of tumors. By combining longitudinal whole genome sequencing (WGS), bulk RNAseq, and single cell (sc)RNAseq spanning plasma and tumor tissue from pet dogs OS we elucidate metastasis-associated genomic and cellular programs within individual patients and nominate biologically grounded, potentially actionable pathways relevant to human OS.

## Results

### Primary and Metastatic OS data is representative of pediatric OS

OS is characterized by a complex, chaotic mutational landscape^7,10,11,13,14,18^. Recent evidence supports conservation of copy number changes in murine models of human OS^14^ and genomic instability fueled by chromothripsis events driving clonal tumor evolution^10^. To ensure our dataset was representative of pediatric OS, we first evaluated the genomic landscape via WGS.

Consistent with published datasets, the landscape of single nucleotide variants (SNVs), structural variants (SVs) and copy number variants (CNVs) was highly diverse among the primary and metastatic tumor samples analyzed by WGS (Figure 1A-B; SFig 1A-B; STable 1; STable 2; STable 3; STable 4), spanning multiple event classes including missense mutations, frameshift mutations, and copy number gains and losses. As observed in previous studies, *TP53* exhibited SNVs and SVs in 82% of all sequenced samples; no copy-number alterations affecting the *TP53* locus were detected (Figure 1A; SFig 1A)^7,8,13,19,20^. Other mutated genes previously reported in OS were similarly altered in our dataset, including copy number gains in *MYC (38%, 22/58)*, copy number loss in *DLG2 (28%, 16/58),* and large deletions in *DMD (22%, 13/58)*^7,8,13^.

**Figure 1:**
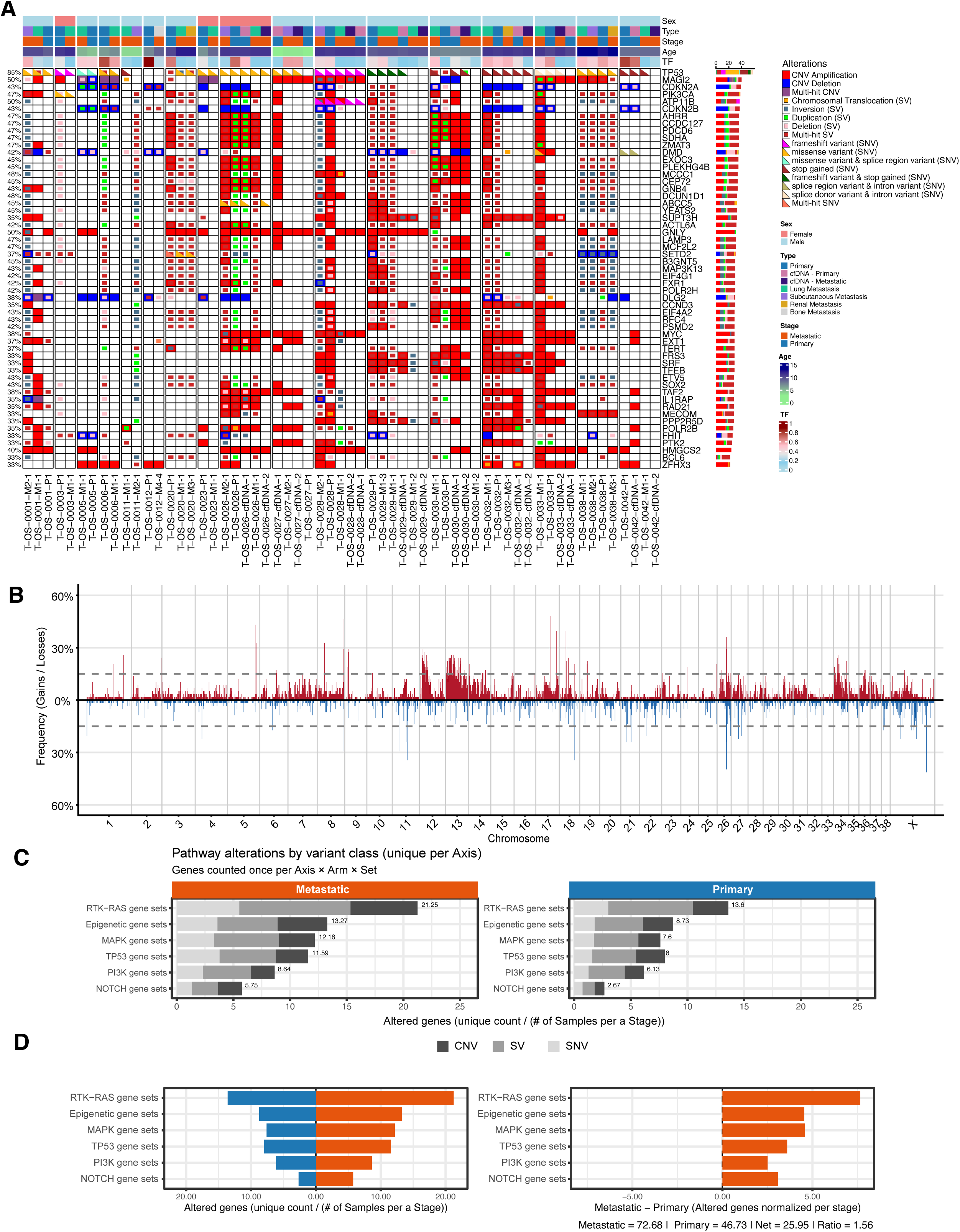
Genomic landscape and mutational burden across primary and metastatic osteosarcoma samples. (**A**) Oncoprint showing mutations present in at least 33% of samples, subsetted to the top 20 mutated genes, COSMIC Tier 1 Consensus genes, and recurrent osteosarcoma-associated pathways. Each column represents an individual dog and includes single-nucleotide variants (SNVs), copy-number variants (CNVs), and structural variants (SVs). Top bars indicate sex, sample type, age, and tumor fraction (TF). **(B)** Frequency of focal (< 3 Mb) copy number gains (red, above axis) and losses (blue, below axis) plotted in 1 Mb bins across all autosomes and chromosome X. Each bar represents the proportion of samples harboring focal alterations at that genomic position. Dashed horizontal lines indicate the 15% recurrence threshold. Recurrent gains are prominent on chromosomes 13 and 31, while recurrent losses are observed on chromosomes 26 and X. **(C)** Stacked bar plots showing the normalized number of altered genes per pathway gene set in metastatic (left, orange) and primary (right, blue) tumors. Genes were counted once per pathway and normalized by the number of samples at each disease stage. Bars are subdivided by variant type—copy-number variants (CNV, dark gray), structural variants (SV, medium gray), and single-nucleotide variants (SNV, light gray). RTK–RAS, epigenetic, TP53, and MAPK gene sets exhibited the highest overall alteration burden, with metastatic tumors showing consistently greater pathway-level alteration frequency across all variant classes. **(D)** Comparison of altered pathway gene sets between metastatic and primary osteosarcoma. Bar plots show the normalized number of altered genes per pathway gene set across canonical oncogenic pathways (RTK–RAS, epigenetic, TP53, MAPK, PI3K, and NOTCH). The left panel displays normalized alteration counts by stage (metastatic = orange; primary = blue), while the right panel shows the difference (Met – Prim), where positive values indicate higher alteration frequency in metastases. Overall, metastatic samples harbored a greater burden of pathway alterations (Met = 72.68; Prim = 46.73; net = 25.95; ratio = 1.56).

In addition to extensive amplification of chromosome 13, where MYC and KIT are located in pet dogs, focal copy number gains and losses were present throughout the genome (Figure 1B; STable 5). The strongest focal gains appeared on chromosomes 17 (48%), 8 (47%), 5 (43%), and 18 (40%), which overlap *MAGI2, NOTCH2, GNLY, KDM3A* and *ZFHX3*. Additional genes amplified on the same chromosomes outside of the focal regions include *NRAS, SETDB1, SLC16A1, RHOC, CHEK1,* and *WWOX*. Recurrent focal losses clustered on chromosomes 35 (53%), 17 (48%), X (41%), 11 (29-34%) and 26 (40%). These regions overlap OS relevant genes, including *CDKN2A* and *CDKN2B*. Chromosome 34, syntenic with human 3q26, experienced recurrent gains of 11–12 Mb (26%).

Chromothripsis events were widespread, affecting 32 of 39 chromosomes. Of the 148 events identified, 82 were found in metastases and 61 in primary tumors (Stable 6; SFig 2A). Chromothripsis was most common on chromosomes 34 and 20, both of which were significantly enriched after adjusting for chromosome length (SFig 2A-B). Recurrent gains on chromosome 34, syntenic with human 3q26, provide the strongest evidence for cross-species conservation, as this region is recurrently altered across multiple human cancers and has been previously implicated in canine OS^21^. The region of chromothripsis on chromosome 20 also overlaps *SETD2* which is recurrently mutated in canine OS^7,13^.

### Recurrently mutated genes demonstrate stage-focused enrichment of immune and DNA damage repair pathways

While hotspot mutations are rare in OS, genomic changes present in our OS samples typically impacted similar pathways such as RAS, PI3K, MAPK, NOTCH, TP53 and DNA damage repair, as well as chromatin modifying genes. Several of these are listed in the CiViC database as being associated with therapeutic resistance^7,10,19,20,22–27^. We evaluated CNV, SV, and SNV events across these pathways to determine whether they were more frequently altered in metastatic lesions compared to the primary tumors (Fig. 1C). Metastatic tumors showed a higher overall number of alterations, especially in the PI3K, MAPK, and epigenetic axes (STable 7; Fig. 1D). Paired CNV analysis revealed nominally significant pathway differences between primary and matched metastases for PI3K (p-adjusted = 0.084), epigenetic (p-adjusted = 0.070), MAPK (p-adjusted = 0.087), NOTCH (p-adjusted =0.087), RAS (p-adjusted = 0.11), and TP53 (p-adjusted=0.11). Sign tests further demonstrated that these differences were directionally consistent, with metastases showing greater CNV gains for PI3K (fraction = 1.00, p = 0.0078; p-adjusted = 0.046), and epigenetic pathways (fraction = 0.875, p = 0.070; p-adjusted 0.21) (STable 8). Together, these results suggest widespread amplification of oncogenic and chromatin-remodeling programs in metastases.

Tumor cells proliferate under a variety of pressures throughout the disease course, including clonal and therapeutic selection of tumor cell subpopulations that dictate the genomic composition of metastatic outgrowth^10^. Consistent with clonal selection during metastatic dissemination, certain alterations were enriched in primary tumors while others were more frequent in metastases, suggesting that metastatic clones derive from subpopulations with distinct genomic profiles. Metastatic tumors more often showed a high number of alterations in *ATRX*, *VEGFA*, *ATR*, ATM, *CCND3*, *PDGFRA*, *BRCA1/2*, *KDM6B*, *PALB2*, and *LRP1B*, indicating reinforcement of MAPK, PI3K, and DNA damage repair pathways (Fig. 2A). Conversely, primary tumors more frequently had changes in *CDKN2A*, *DMD*, *PTEN*, *BAP1*, *MAP2K4*, *KMT2D*, *PDGFRB*, *DNMT3A*, *TP53*, and *DLG2*, reflecting loss of genetically altered genes in cells susceptible to therapeutic treatments (Fig. 2A). When we cross-referenced genes mutated exclusively in metastatic samples with the CiViC database, 17 overlapped, including *MTOR*, *BRAF1*, *EGF*, *FGF2*, *CTNNB1*, *AURKA*, *PROM1* (CD133), *SIRT1*, *NQO1*, *PBK*, and *GNAS*, which are implicated in OS biology^28–38^. Furthermore, six genes overlapped with the CiViC database, including *CD274* (PD-L1), *BAP1*, *DNMT1*, *JAK2*, and *RET*, which have been previously linked to immune evasion and growth control in OS (STable 9)^39–42^.

**Figure 2.**
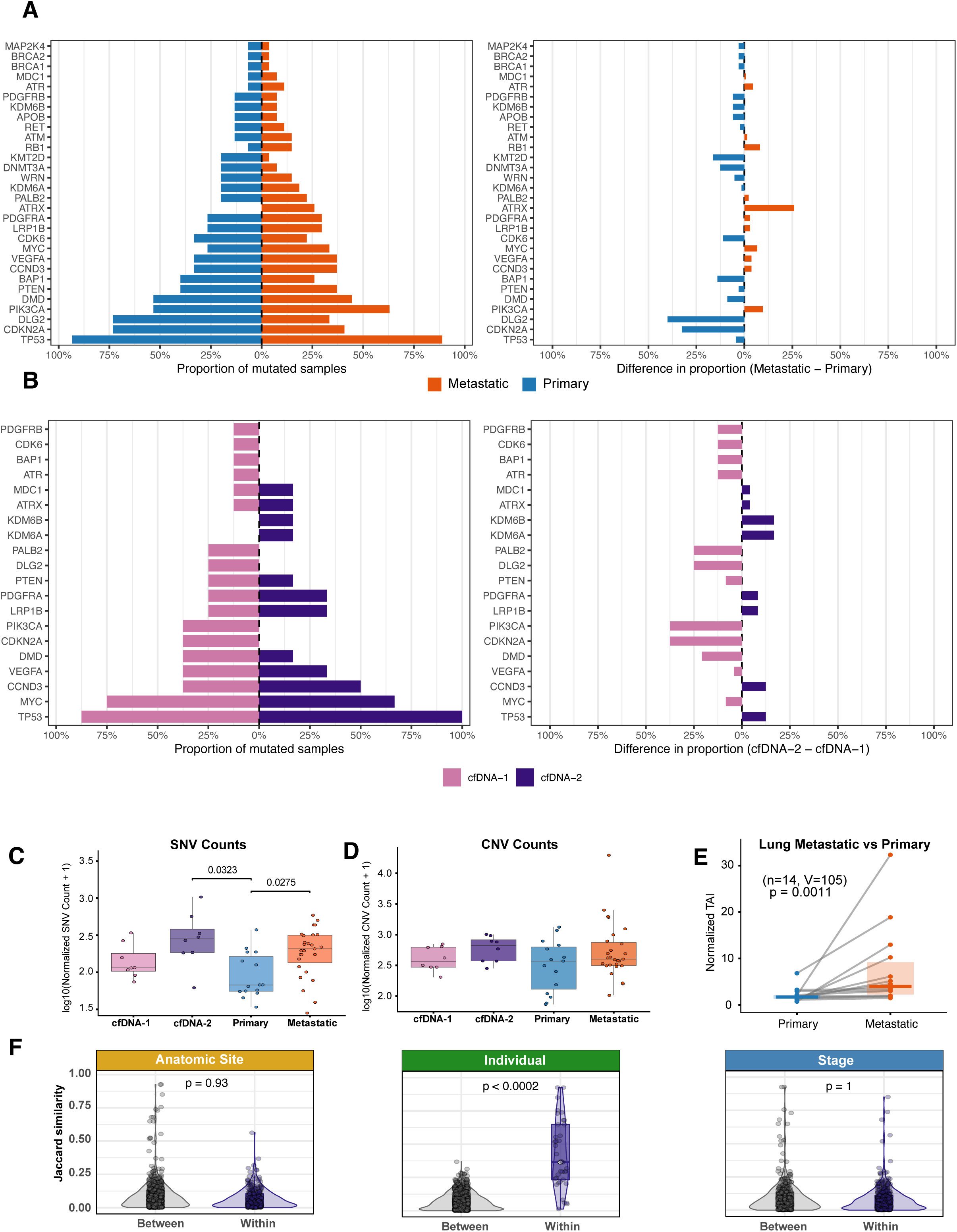
Oncogenic pathway alterations by variant class and disease stage in osteosarcoma. **(A)** Frequency of somatic mutations in cancer-associated genes comparing primary tumors (blue) and metastatic samples (orange). Left panel shows the proportion of samples harboring mutations in each gene. Right panel shows the difference in mutation frequency (metastatic − primary), where positive values (orange) indicate enrichment in metastases and negative values (blue) indicate enrichment in primary tumors. Genes are ordered by total mutation frequency. **(B)** Frequency of somatic mutations detected in longitudinal circulating free DNA (cfDNA) samples. cfDNA-1 (pink) represents the initial collection timepoint and cfDNA-2 (purple) represents a subsequent timepoint. Right panel shows the difference in mutation frequency (cfDNA-2 − cfDNA-1), where positive values indicate increased detection at the later timepoint. **(C)** Boxplots of function-impacting SNVs normalized by tumor fraction for each sample. Metastatic tumors exhibited a significantly higher burden of mutations compared to primary tumors (p_adj = 0.0275). cfDNA-2 samples also differed significantly from primary tumors (p_adj = 0.0323). **(D)** Boxplots of normalized CNV counts showing no significant differences between sample types, although cfDNA and tissue samples exhibited distinct distributions. **(E)** Genomic instability, measured as normalized total aberration index (TAI), was higher in lung metastases than in matched primary tumors (p = 0.0097). **(F)** Violin plots showing Jaccard similarity of shared mutations within and between anatomical sites, individuals, and stages. Samples from the same individual were significantly more similar than those between individuals (p < 0.0002), while no significant differences were observed by site or stage.

Among metastasis-specific SNVs, we observed a significant enrichment for EGF-like domains, including *LRP1B* and *NOTCH3/4* (FDR=0.0035; STable 10). Pathway enrichment of metastasis-specific CNVs revealed enrichment for immune and inflammatory signaling, including IL-1 receptor binding (p-adjusted= 4.7 × 10⁻⁸), NF-κB and Toll-like receptor pathways (p-adjusted=0.005), and cytokine-receptor binding (p-adjusted = 0.0017; STable 11). These findings indicate that metastatic progression of OS involves remodeling of the tumor immune microenvironment.

Genomic instability was also reflected in frequent DDR pathway alterations across all samples, particularly affecting double-strand break repair (HR, NHEJ) and checkpoint genes such as *ATR, ATM, BRCA1/2, PALB2, BAP1*, and *DLG2*. Notably, while primary tumors showed a trend toward higher SV burden, metastases exhibited significantly elevated SNV burden in DDR genes (paired Wilcoxon p = 0.008; STable 12), suggesting a shift in mutational mechanisms during disease progression. Lung metastases in particular showed elevated total aberration index (TAI) compared to matched primaries, consistent with increased chromosomal instability at metastatic sites. Across all samples, DDR genes were frequently altered, with SVs accounting for 39.6% (146/369) of events, followed by SNVs (32.0%, 118/369) and CNVs (28.5%, 105/369). In primary tumors, SVs accounted for 45.3% (48/106), SNVs 28.3% (30/106), and CNVs 26.4% (28/106). In metastases, SVs represented 37.3% (98/263), SNVs 33.5% (88/263), and CNVs 29.3% (77/263) (STable 12) with additional SV and SNV contributions, especially in double-strand break repair (HR, NHEJ) and checkpoint genes such as *ATR*, *ATM*, *BRCA1/2*, *PALB2*, *BAP1*, and *DLG2*. To compare DDR burden between stages, we calculated per-sample DDR alterations in matched primary-metastasis pairs. Primary tumors had an average of 32.2 SVs, 13.7 SNVs, and 14.2 CNVs affecting DDR genes per sample, while matched metastases showed 23.8 SVs, 17.0 SNVs, and 17.5 CNVs (paired Wilcoxon signed-rank test: SV p = 0.065, SNV p = 0.008, CNV p = 0.514; n = 12-14 pairs). Notably, while primary tumors showed a higher SV burden, metastases exhibited significantly elevated SNV burden in DDR genes, suggesting a shift in the mutational mechanisms affecting DNA repair pathways during disease progression. These findings highlight the role of genome instability and replication stress in advanced treatment-resistant stages of OS.

### Patient identity is the dominant organizer of the WGS mutational landscape

Given the relatively limited patient-matched WGS datasets available in people^18^, understanding how OS evolves within individual patients remains challenging. We leveraged the conserved OS mutation landscape in pet dogs to characterize intra-patient heterogeneity across early and late stage patient-matched OS samples. Comparing mutational burden, copy number architecture, and chromothripsis patterns across paired primary and metastatic tumors enabled us to identify genomic features associated with disease progression (Figure 2C-F).

Across the cohort, copy number changes did not significantly differ between metastatic tissue sites. Stratification of samples by tumor stage revealed distinct copy number patterns associated with metastasis (SFig 1B). Lung metastases exhibited chrX losses (5.1-fold, p < 10⁻⁴³), chr11 gains (4.4-fold, p < 10⁻²²), and chr12 losses (6.9-fold, p < 10⁻²¹) when compared to primary tumors. Genome-wide instability was significantly higher in lung metastases compared to primary tumors (p = 0.0091; Figure 2E), however this was not attributable to an increase in chromothripsis. Of the 15 patients with matched primary and metastatic samples, 10 exhibited chromothripsis in at least one sample. Among these 10 patients, 70% (7/10 patients) shared at least one chromothripsis-affected chromosome, indicating stable patterns of chromothripsis within individual patients.

Mutational burden was significantly higher in metastatic lesions compared to primary tumors. Metastatic samples had more mutations than primary samples among functional variants (p=0.0275; Figure 2C; SFig 2C; STable 13). This difference remained significant in paired within-patient analysis, demonstrating that individual patients accumulate mutations during disease progression (n = 24 pairs; p = 0.027). When all patient-matched samples were considered together, lung metastases had significantly higher mutation burdens than primary tumors (SFig 2D-E, STable 14). However, significance was lost when evaluated within paired patient-specific samples, likely due to reduced statistical power (n = 17 pairs; p = 0.10).

To better characterize the influence of patient identity versus tumor stage on mutational landscape, we evaluated mutations in the context of which genes were impacted, rather than considering each mutation type and count separately. When evaluated in this manner, samples from the same patient were significantly more similar across structural, copy-number, and single-nucleotide mutations than samples from different patients (p < 0.0002; Fig. 2F), a finding further supported by ECDF analysis demonstrating statistically distinct distributions across patients (p < 0.001; SFig. 3A-B). Anatomic site and disease stage showed no significant effects (p = 0.33 and p = 0.089, respectively; STable 15). Sensitivity analyses to ensure our results were not being skewed by bone and renal metastases (which had a relatively low sample size) did not identify an impact (SFig. 4A-B; STable 15). Consistent with these pairwise analyses, we confirmed that patient identity is the most significant predictor of mutational composition across all datasets, explaining 49–60% of variance (p = 0.0002; SFig 5A; STable 16). In contrast, anatomic site accounted for only 2–6% of variance (p > 0.05), and age had minimal impact (p > 0.17; SFig. 5A; STable 16). Because recurrently mutated genes may represent biologically significant drivers in OS, we assessed whether patient identity continued to dominate mutational similarity within this subset. Unsupervised hierarchical clustering of Jaccard similarity scores for OS driver genes also grouped samples by patient rather than disease stage, reinforcing patient identity as the dominant determinant of mutational composition (SFig. 3C).

### cfDNA Recapitulates Stage-Matched Tumor Mutation Profiles

Liquid biopsy is a powerful approach for identification of tumor-relevant genetic variants^43–45^. In OS, high level copy number alterations are typically noted, providing valuable structural information for characterization of tumor fraction and minimal residual disease monitoring. Evaluation of (30X) WGS of patient and stage matched cfDNA alongside tumor tissue enabled us to characterize stage-specific tumor evolution of the genomic variant landscape in cfDNA within individual patients.

SVs were detected in cfDNA, however when compared to dog-matched tumor tissue the frequency of calls was diminished. SV detection in cfDNA is associated with challenges inherent to the fragmented nature of cfDNA, resulting in breakpoints located in fragmented regions of cfDNA and relatively lower levels of tumor purity in these samples that are best overcome with hybrid capture approaches that enable significantly higher coverage over these regions compared to standard WGS. To ensure that the difference in SV detection did not influence our conclusions, we did not evaluate SVs when comparing mutation profiles across tissue and plasma, and instead focused on SNV and CNV detection^46,47^.

Consistent with tumor tissue, cfDNA collected at the time of metastasis (cfDNA-2) showed higher overall mutation burden compared to cfDNA collection at the time of diagnosis of the primary tumor (cfDNA-1) (function-impacting variants p = 0.032; all non-synonymous variants p = 0.013) (Figure 2C, SFig 2C, STable 13), with similar effect sizes in patient-matched paired samples (δ = 0.53). Beyond total burden, the mutational composition of cfDNA-2 shifted toward a metastatic profile, with a gain of metastasis-specific variants (paired OR = 9.33; p = 4.65×10⁻⁶), while primary-specific and shared variants were lost (OR = 0.077 and 0.224, both p < 0.02). Jaccard similarity analyses confirmed this shift (ΔJaccard p = 0.007; permutation p = 0.0039, SFig 6A). Together, these results indicate cfDNA variants reflect both the burden and clonal composition of stage-matched tumor tissue.

Mutations observed in tissue-identified genes are also captured in cfDNA. Compared to cfDNA collected at baseline (cfDNA-1), cfDNA-2 had numerically higher mutation frequencies in *CCND3, KDM6A, KDM6B, LRP1B, VEGFA, TP53, ATRX,* and *MDC1*, paralleling metastatic profiles. However, cfDNA-1 retained higher levels of *ATR, BAP1, DMD, PIK3CA, PTEN, DLG2, CDK6, PALB2, PDGFRB*, and *MYC*. These longitudinal differences imply that metastatic cfDNA preferentially captures mutations in signaling and chromatin-remodeling drivers. In contrast, pre-treatment cfDNA predominantly reflects baseline DNA-repair deficiencies and innate tumor suppressor gene loss. Together, cfDNA and tissue provide complementary views of mutational processes. They support a combined approach where cfDNA increases sensitivity to focal events, and tissue analysis reveals large-scale structural features that inform stage-specific features and potential therapeutic opportunities.

Copy number alterations in cfDNA also largely reflected those observed in matched tissue samples. The overall number of copy number changes did not differ significantly between cfDNA and tissue (Figure 2D; SFig. 6B). cfDNA recapitulated key tissue copy number patterns, including gains on chr13 and losses on chrX, which harbor *KIT, MYC*, and *DMD*. However, genome-wide instability was lower in cfDNA-2 compared to lung metastases (p = 3.45e-2; SFig 6C), likely reflecting the lower tumor fraction and fragmented nature of cfDNA. No other site comparisons reached significance.

### Inter-patient variability dominates both cell type composition and transcriptional profiles

To better understand the impact of inter-patient genomic heterogeneity on transcriptional signature dynamics, we leveraged bulk and sc-RNA sequencing to characterize the dynamics of gene expression in paired treatment naïve and advanced stage OS samples. Graph-based clustering and UMAP projection of scRNAseq from five dogs spanning 12 primary and metastatic OS samples (41,297 cells) revealed 13 transcriptionally distinct populations of nuclei grouped into four main compartments: tumor (malignant osteoblasts), myeloid, lymphoid, and stromal (Fig. 3A–B). Malignant osteoblasts were abundant in several primary tumors, and nuclei from individual patients clustered as different subtypes, underscoring the strong inter-patient heterogeneity characteristic of the tumor mutational landscape. Immune populations (T, B, and myeloid cells) were less common and varied widely across individuals. Tumor and stromal cells comprised the majority of cells at both primary and metastatic sites. There was no consistent trend in the ratio of tumor and stromal cells across samples (Fig. 3C). Most metastatic samples (6/8 samples) contained abundant malignant osteoblasts, whereas two were enriched for fibroblasts. The tumor and stromal compartments dominated both primary and metastatic OS samples, but their relative proportions, especially of malignant osteoblasts and fibroblasts, varied markedly between individuals.

**Figure 3.**
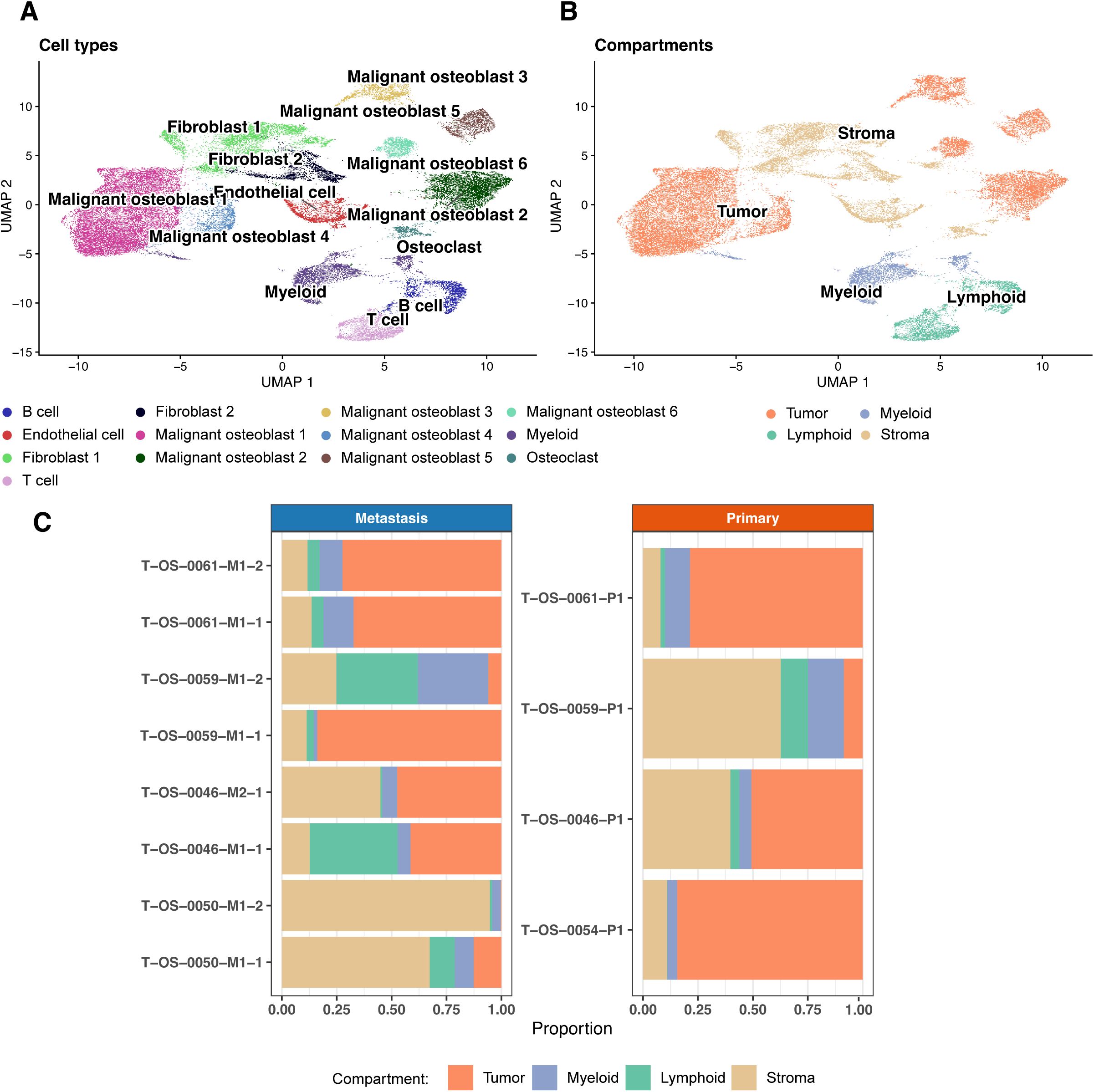
Cellular composition of primary and metastatic canine osteosarcoma by by single-cell-resolution transcriptomics. **(A)** UMAP visualization of all cells colored by annotated cell type, including malignant osteoblast subtypes (1–6), fibroblasts, myeloid cells, lymphocytes (B and T cells), osteoclasts, and endothelial cells. Malignant osteoblast clusters exhibit transcriptional heterogeneity, separating into six distinct subtypes. **(B)** UMAP visualization of the same dataset grouped into four major compartments: tumor, myeloid, lymphoid, and stroma. These categories encompass malignant, immune, and stromal populations within the osteosarcoma microenvironment. **(C)** Bar plots showing the proportional composition of cell compartments within matched metastatic (left, blue) and primary (right, orange) tumors from individual dogs. Tumor compartments dominated both stages, but metastatic samples displayed greater heterogeneity in myeloid and stromal cell representation compared to their matched primary tumors.

We characterized variability in gene expression based on OS stage, patient, and cell type composition. Expression levels were more similar within than between patients for the 2000 most variably expressed genes using metrics designed to evaluate concordance of ranked variables and gene expression abundance (Spearman p = 0.001; Bray-Curtis p = 0.002) (Fig. 4A; SFig. 7A). Disease stage showed no significant difference, further supporting that tumor composition is largely patient-specific (Spearman p = 0.887; Bray-Curtis p = 0.779) (Fig. 4B; STable 17). Composition-based similarities displayed an even larger within-patient shift (Fig. 4A; STable 17) (Spearman p = 0.022; Bray–Curtis p = 0.005), while stage effects remained weak or absent (Fig. 4B; STable 17) (Spearman p = 0.960; Bray–Curtis p = 0.920). Both analyses showed a rightward shift in within-patient similarity compared to between-patient comparisons, confirmed by Kolmogorov–Smirnov (KS) tests (Bray–Curtis p = 0.0024; Spearman p = 0.0065), indicating that patient identity is the main factor influencing similarity even when combining compositional and expression-based data (SFig. 7B-C).

**Figure 4.**
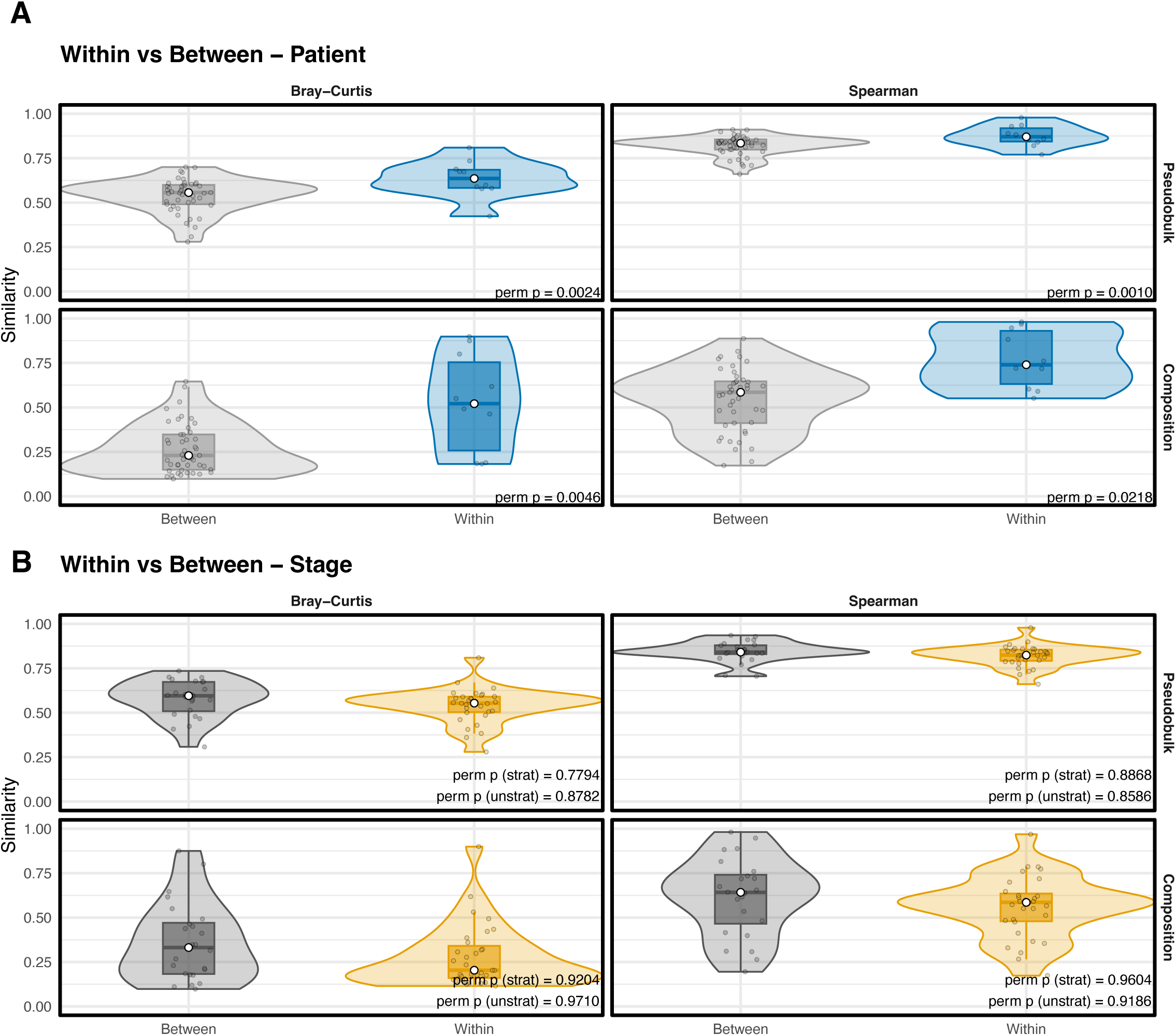
Intrapatient similarity exceeds interpatient variation in tumor composition and pseudobulk expression. **(A)** Violin plots showing within- versus between-patient similarity in both pseudobulk gene expression (top) and cell-type composition (bottom). Similarity was measured using Bray–Curtis dissimilarity (left) and Spearman correlation (right). Samples from the same patient (within) were significantly more similar than those from different patients (between) across both metrics (Bray–Curtis p = 0.0024; Spearman p = 0.0010 for pseudobulk; Bray–Curtis p = 0.0046; Spearman p = 0.0218 for composition). **(B)** Violin plots showing within- versus between-stage similarity (primary vs. metastatic) using the same distance metrics. No significant differences were observed for either pseudobulk profiles or cell-type composition (all p > 0.05), indicating that stage contributes less to overall transcriptional and compositional similarity than patient identity.

To quantify these effects, we asked how much of the variance in the expression profiles can be attributed to patient identity, disease stage, or age (Fig. S5B; STable 18). Patient identity accounted for the largest share of variance (R² = 0.45–0.47, p = 0.008–0.017), confirming a strong patient-specific signal. In contrast, disease stage and age contributed minimal additional variance (Stage R² = 0.07, p = 0.42; Age R² = 0.03–0.05, p = 0.62–0.80). These results indicate that differences between patients primarily shape the transcriptional landscape, with age and stage playing comparatively minor roles.

### Tumor and Stromal Compartments Dominate the Osteosarcoma Landscape, Revealing Distinct Malignant Osteoblast States

Emerging data suggests that tumor and stromal cells exhibit unique phenotypes in the context of the tumor microenvironment^48–51^. Pathway analysis of non-tumor stromal compartments identified co-localization of immune signaling and tumor microenvironment remodeling, emphasizing coordinated regulation of non-malignant compartments. Conversely, most malignant osteoblast (MOB) clusters were transcriptionally distinct, often forming clusters with other MOB subsets. Examination of the top five marker genes for each population outlined clear transcriptional boundaries, confirming strong cluster identities and high intra-cluster coherence. Non-malignant populations, such as immune, endothelial, fibroblast, and osteoclast, remained well separated, supporting accurate cell-type annotation and highlighting the distinct transcriptional structure of the tumor microenvironment.

Malignant osteoblast (MOB) subclusters exhibited some overlapping gene expression programs, indicating shared features of proliferation, metabolism, and osteomimicry ^52^ (SFig. 8A). MOB3 and MOB5 clustered together due to shared proliferative and metabolic traits, while MOB1 and MOB4 aligned based on stress and immune-related signatures. MOB2 and osteoclasts clustered together, suggesting activation of bone-remodeling pathways. MOB6 showed transcriptional overlap with proliferative clusters (MOB3/5) but was uniquely enriched for mitotic spindle signaling, indicating strong engagement of the cell-division machinery and increased mitotic activity. Clustering of top marker genes across lineages (SFig. 8B; STable 19) confirmed that most non-malignant populations (immune, endothelial, fibroblast, and osteoclast) remained distinct, while malignant osteoblast subtypes showed partial convergence. Pathway enrichment analysis (SFig. 9A; STable20; STable21) revealed all subclusters were enriched for core proliferative and metabolic pathways (E2F targets, G2M checkpoint, MYC targets, oxidative phosphorylation), reflecting a shared malignant basis driven by cell-cycle and energy demands. Stress and immune signaling pathways (TNFα/NF-κB and the interferon-γ response) were also broadly activated, supporting a common inflammatory state. Additionally, subtype-specific adaptations emerged: MOB1 showed reduced PI3K signaling, suggesting decreased growth factor dependence and increased reliance on stress and immune pathways ^53^; MOB2 upregulated fatty-acid metabolism, glycolysis, and peroxisome pathways, indicating lipid-based energy production and an osteoclast-like remodeling phenotype; MOB3 and MOB5 maintained high proliferative programs but diverged metabolically. MOB3 emphasized oxidative phosphorylation and translation, while MOB5 favored glycolysis, mTORC1, and hypoxia signaling, consistent with a stress-adapted proliferator. MOB4 was enriched for angiogenesis, ECM remodeling, and inflammatory modules but uniquely showed suppression of the p53 pathway, implying tolerance to microenvironmental stress. All of the malignant osteoblast (MOB) subtypes were observed in both primary and metastatic samples, suggesting that these transcriptional states are not exclusive to one stage (SFig. 9B). These findings define a continuum of malignant osteoblast states: proliferative (MOB3/5/6), stress-responsive (MOB1/4), and osteomimetic/metabolically specialized (MOB2). Although interconnected by cell-cycle and metabolic activation, subtype-specific pathways reveal distinct adaptive strategies influencing tumor behavior and interactions within the microenvironment.

### Cell–type–resolved pseudobulk expression reveals strong patient specificity in malignant osteoblasts

While tumor cells exhibit high transcriptional variability across patients, we wanted to determine if immune cell phenotypes were conserved. To demonstrate transcriptional organization across different cell types, we generated pseudobulk profiles by averaging expression levels for each individual–cell type pair (Figure 5A). Hierarchical clustering revealed a clear distinction: malignant osteoblasts mostly cluster by patient, whereas tumor microenvironment cells grouped by lineage across individuals. Myeloid, T, and B cells form separate, lineage-specific clusters regardless of the patient, indicating that conserved immune programs shape their transcriptomes. Conversely, malignant osteoblasts from the same patient consistently cluster together and separately from those of other patients (Fisher’s exact p < 0.001; STable 22), highlighting notable inter-patient differences in tumor-intrinsic states that have minimal overlap with other states (Figure 5B–C). These findings suggest patient-specific transcriptional programs primarily drive malignant osteoblast identity, while the non-malignant microenvironment exhibits conserved, lineage-based expression patterns^54^.

**Figure 5.**
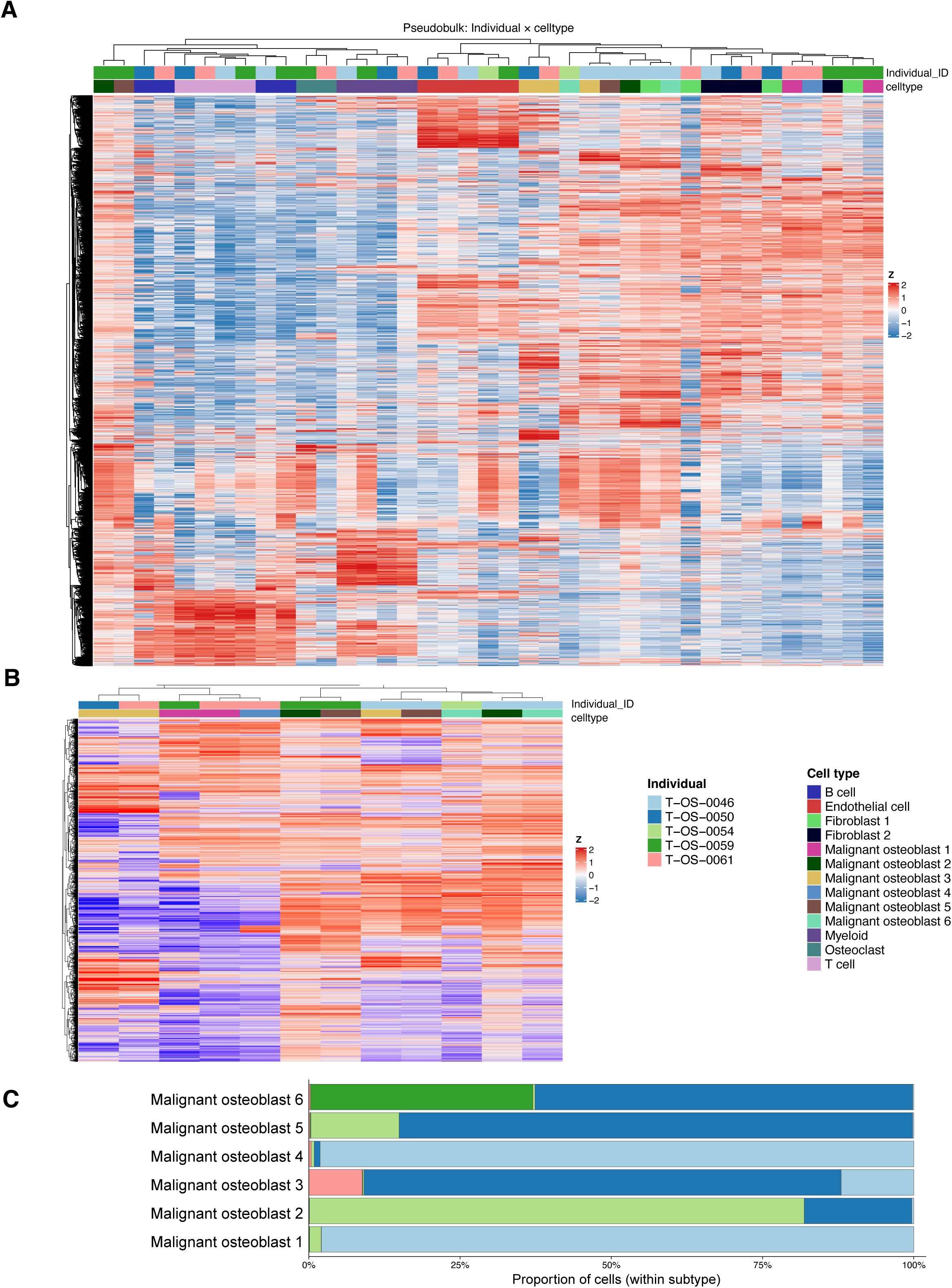
Pseudobulk transcriptional profiles reveal patient-specific clustering and dominant malignant osteoblast subtypes. **(A)** Hierarchical clustering heatmap of pseudobulk gene expression across all major cell types, aggregated by individual. Rows represent genes and columns represent individual × cell type combinations. Expression values are Z-score normalized across samples. Clustering reveals strong patient-specific expression signatures, with samples from the same individual grouping together across multiple cell types. **(B)** Hierarchical clustering heatmap of pseudobulk profiles restricted to malignant osteoblasts, highlighting inter-patient variability and the presence of distinct transcriptional programs among malignant subtypes. (**C)** Stacked bar plots showing the proportional distribution of malignant osteoblast subtypes for each individual. Certain patients exhibit dominance of specific subtypes, suggesting heterogeneity in malignant cell states between individuals.

### Malignant osteoblast subtypes with increased immune infiltration dominate metastatic osteosarcoma lesions

Because the scRNAseq datasets were generated from dogs without WGS data, we used scRNAseq data as a reference for deconvolution of bulk RNA-seq data generated from the same tumor samples used for WGS. This analysis allowed us to validate our conclusions at the single cell level in a larger bulk RNA-seq dataset. Deconvolution of bulk RNA-seq data revealed that tumors are primarily composed of three main malignant osteoblast (MOB) populations (MOB5, MOB2, and MOB6) which together account for most gene expression in both primary and metastatic lesions (STable 23). Other malignant osteoblast states, such as MOB1 and MOB4, were detected only sporadically and at low levels, while MOB3 was not observed in the bulk RNAseq dataset. Among non-malignant stromal cells, fibroblasts and endothelial cells were consistently present at varying proportions across patients (Figure 6A; STable 23). Both non-malignant fibroblast subsets were identified in the bulk RNA-seq cohort. Furthermore, endothelial cell levels were inconsistent across tumors, suggesting variability in the extent of the vascular compartment. Osteoclasts were rare and appeared sporadically. Immune cells, including T cells, B cells, and myeloid cells, contributed less than 10% of the total expression but varied between patients, with increased immune infiltration seen mostly in metastatic lesions. This pattern suggested a higher immune infiltrate in metastases than in matched primary tumors (Figure 6B). While trends in single-cell data indicated immune cell enrichment, associations with stage in bulk RNAseq were not statistically significant (p > 0.05; Figure 6C).

**Figure 6.**
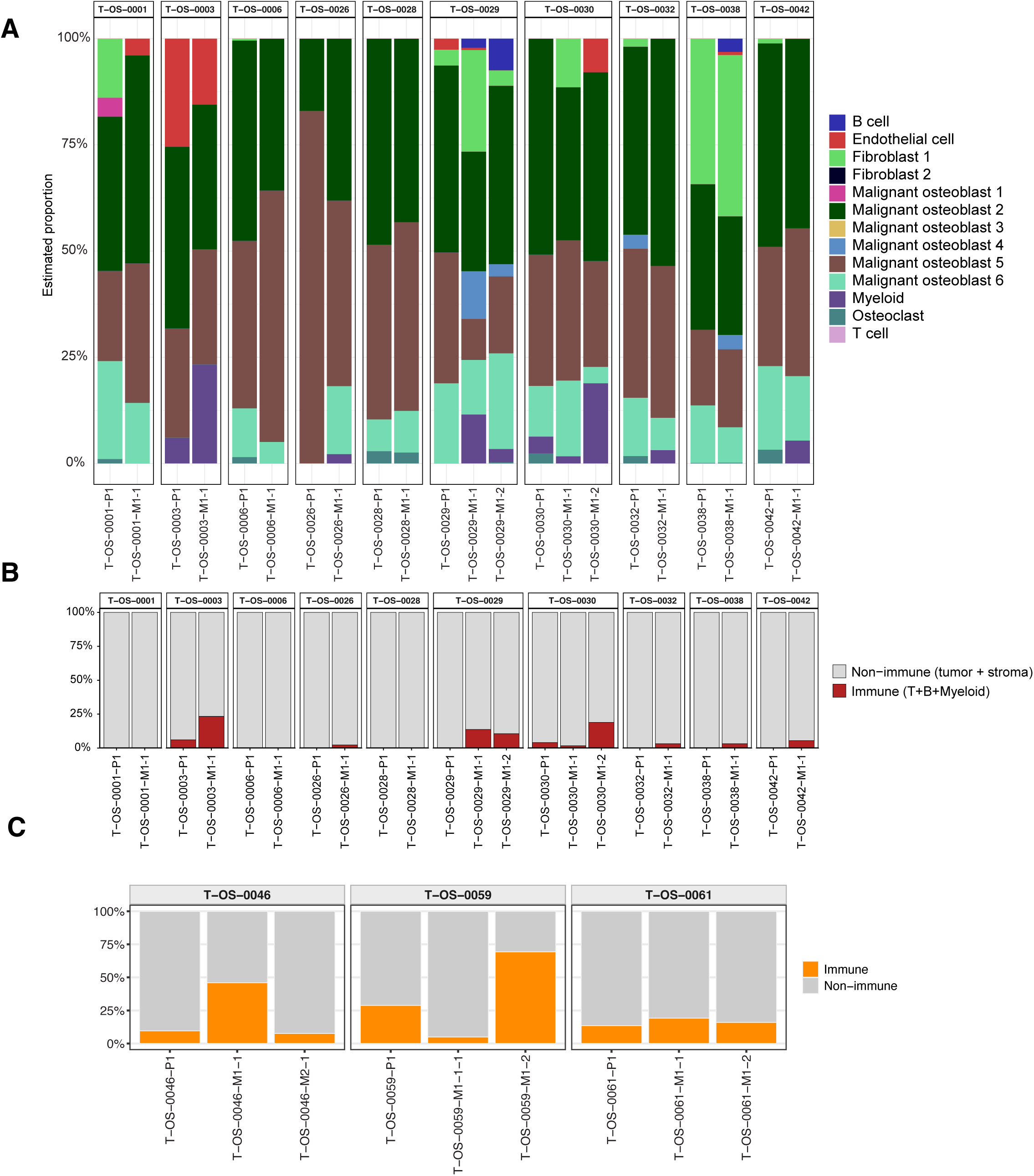
Cell-type composition and immune infiltration across matched primary and metastatic osteosarcoma samples. **(A)** Stacked bar plots showing the proportional composition of major cell types for each patient across matched cfDNA, primary, and metastatic samples using MuSiC deconvolution output. Bars represent the relative abundance of annotated cell populations, including malignant osteoblast subtypes (1–6), fibroblasts, myeloid cells, lymphocytes, endothelial cells, and osteoclasts. **(B)** Aggregated view summarizing the proportion of immune versus non-immune compartments (tumor + stromal) per individual. Although tumor and stromal compartments dominated across most samples, immune infiltration was detected at varying levels between individuals. **(C)** Comparison of immune versus non-immune proportions in matched primary and metastatic tumors for three individuals, showing variable immune composition across disease stages and metastatic sites.

### Immune activation and extracellular remodeling are prominent features of metastases, contrasting with immune-cold malignant osteoblast programs

Functional classification of transcriptional programs differentially expressed in paired primary and pulmonary metastatic osteosarcoma lesions identified 293 differentially expressed genes including 236 upregulated and 57 downregulated in metastases (SFig. 10A–B; STable 24). Metastases were notably enriched for immune and inflammatory pathways, such as INFLAMMATORY RESPONSE, IL6–JAK–STAT3 SIGNALING, COMPLEMENT CASCADE, and ANTIMICROBIAL PEPTIDES, indicating widespread immune activation at the tissue level (STable 25). Simultaneously, pathways such as HYPOXIA, KERATINIZATION, and SURFACTANT METABOLISM were enriched, suggesting active extracellular remodeling and stress responses in the metastatic microenvironment. Conversely, pathways associated with proliferation and metabolism, including OXIDATIVE PHOSPHORYLATION, E2F TARGETS, DNA REPLICATION, BASE EXCISION REPAIR, and DNA METHYLATION, were downregulated, implying that metastatic tumors are less driven by cell-cycle activity and more influenced by microenvironmental signaling. This highlights a transition from proliferative to immune–stromal-dominant transcription in metastatic OS.

We modeled the immune fraction, which is the combined proportion of T cell, B cell, and myeloid cells relative to disease stage, using a mixed-effects model that accounted for patient-level dependence. This analysis showed significantly higher immune representation in metastases compared to primary tumors (odds ratio = 324.6, p = 0.0039) (Fig. 6B, Fig. 7A; STable 26). A sensitivity analysis using patient-level medians for each stage confirmed this result (paired Wilcoxon test, P = 0.011), reinforcing that immune enrichment in metastases is consistent and not due to pseudo-replication within patients.

**Figure 7.**
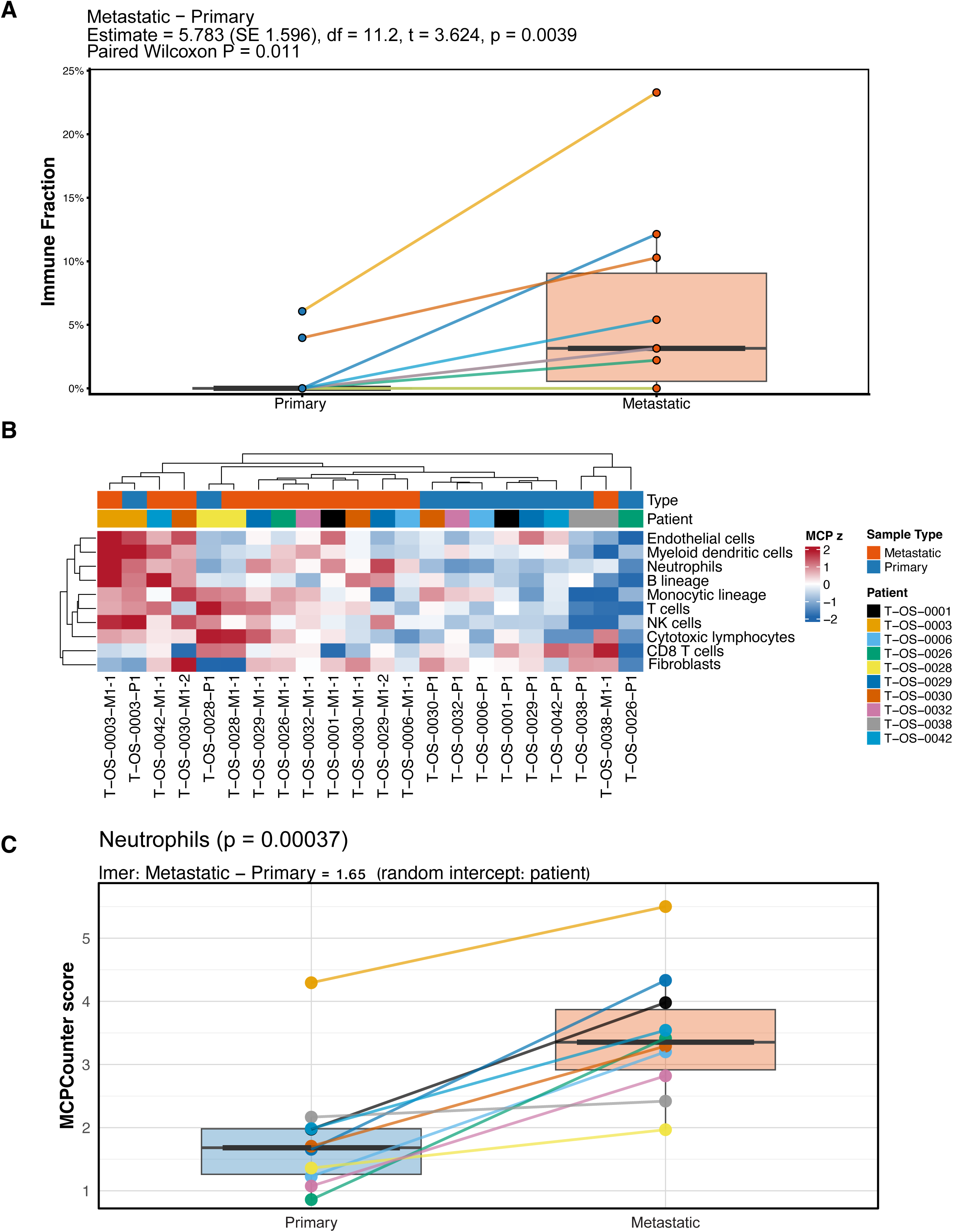
Increased immune infiltration and neutrophil abundance in metastatic osteosarcoma. **(A)** Boxplot showing the overall immune fraction in paired primary and metastatic tumors from the same individuals. Each line connects matched samples, illustrating a significant increase in immune fraction in metastases (mixed-effects p=0.0039; *p* = 0.011, paired Wilcoxon test). **(B)** Heatmap of MCPcounter scores for individual immune and stromal populations across matched primary and metastatic samples. Rows represent cell types, and columns represent samples. Higher scores (red) indicate greater estimated abundance. Several immune subsets, including neutrophils and monocytic lineages, showed increased representation in metastatic tumors.**(C)** Boxplot of MCPcounter-derived neutrophil scores, showing significantly higher neutrophil abundance in metastatic compared to primary tumors (*p* = 0.00037; linear mixed-effects model, random intercept = patient).

Composition-aware analyses further supported this pattern, identifying significantly higher levels of immune and other stromal cells in the metastatic tumor microenvironment that clustered together in hierarchical analysis (Fig. 7B). Moreover, we observed increased neutrophil abundance in metastases (estimate = 1.65; p-adj = 0.00037; STable 27; Fig. 7C), while single-cell deconvolution predicted higher proportions of immune cells (T cell, B cell, and myeloid). Consistent with this, scRNA-seq revealed that malignant osteoblast (MOB) populations are intrinsically immune-cold but metabolically active, characterized by upregulation of OXIDATIVE PHOSPHORYLATION, MYC, E2F, and DNA REPAIR programs, alongside suppressed cytokine and inflammatory pathways (IFN-γ RESPONSE, TNFA/NF-κB, and IL6–JAK–STAT3). In contrast, immune cell clusters exhibited strong enrichment for inflammatory signaling, mirroring the bulk-level observations (SFig. 8A). Collectively, these findings indicate that the immune activation observed in metastatic OS arises from increased immune and stromal infiltration rather than intrinsic tumor reprogramming, underscoring stage-dependent remodeling of the metastatic microenvironment.

### Distinct roles of patient identity and disease stage in shaping tumor and immune transcriptional profiles

We compared similarities within and between groups using normalized expression data and cell-type compositions to determine whether transcriptional similarity among bulk RNA-seq samples is primarily influenced by patient identity or disease stage. For overall expression, similarity within the same patient consistently surpassed that between different patients across both metrics (Bray–Curtis p = 0. 0002; Spearman p = 0. 0002) (Fig. 8A; SFig. 11A-B; STable 28), signifying that transcriptomes from the same individual are significantly more similar than those from different patients. This consistent separation across both magnitude-and rank-based metrics highlights the dominance of strong, patient-specific transcriptional programs. In contrast, the influence of disease stage was less pronounced and more variable (Fig. 8B). Primary and metastatic samples largely overlapped, with only modest separation identifiable via Spearman correlation (stratified p = 0. 0022) (STable 28), and no significant difference was observed with Bray–Curtis (stratified p = 0. 25).

**Figure 8.**
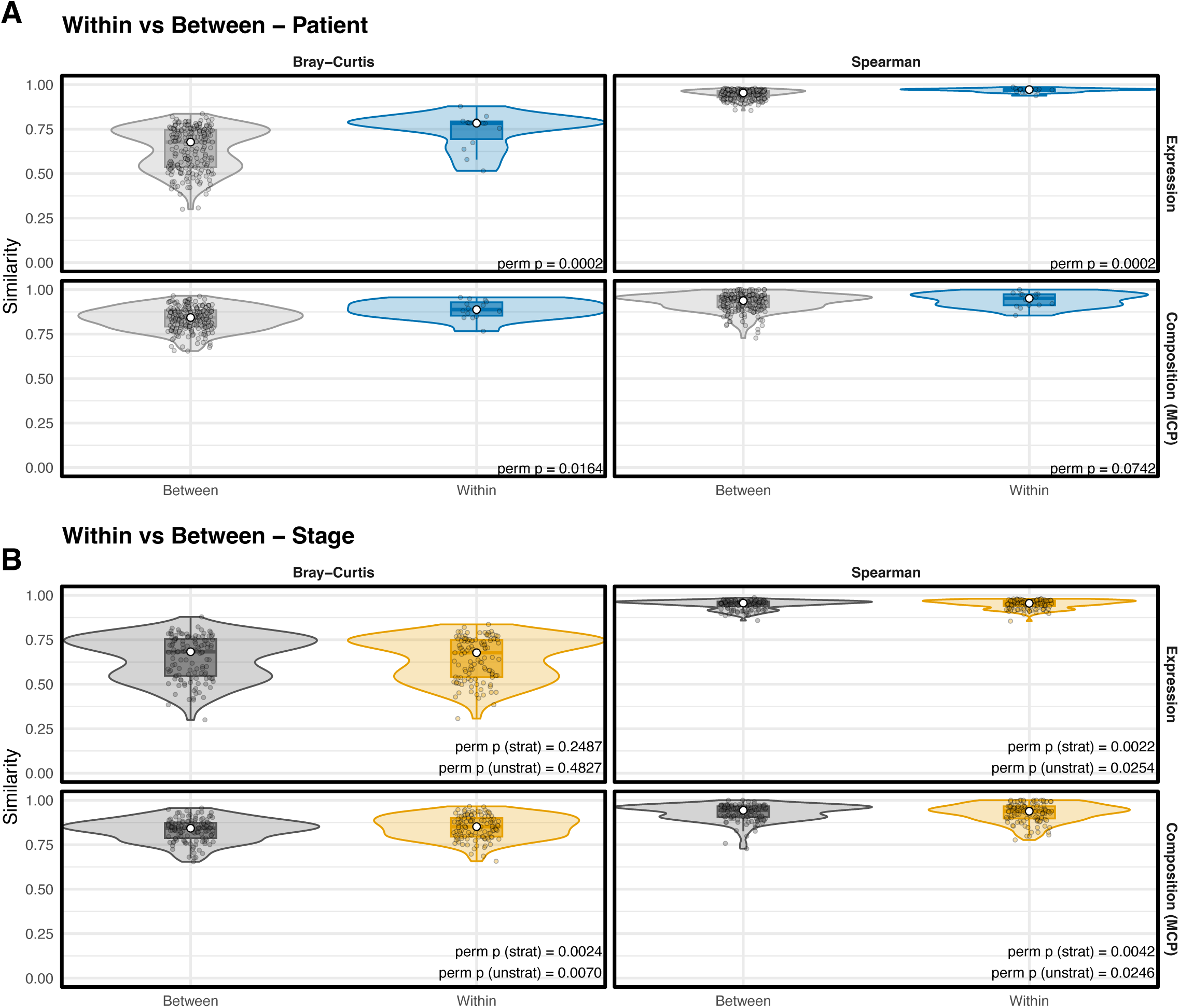
Intrapatient similarity in immune cell expression and composition exceeds interpatient and stage-specific variation. **(A)** Violin plots showing within- versus between-patient similarity for both expression (top) and MCPCounter output (bottom). Similarity was calculated using Bray–Curtis dissimilarity (left) and Spearman correlation (right). Samples from the same patient (within) were significantly more similar than those from different patients (between) for expression (Bray–Curtis p=0.0002; Spearman p=0.0002) and composition by Bray–Curtis (p=0.0164), though Spearman correlation for composition showed a similar trend that did not reach significance (p=0.0742). **(B)** Violin plots comparing within- versus between-stage similarity. Within-stage samples showed significantly higher similarity for composition (Bray-Curtis p=0.0024; Spearman p=0.0042) and expression (Spearman p=0.0022), though Bray-Curtis expression similarity did not differ significantly by stage (p=0.2487).

Conversely, composition-based similarities exhibited an opposite pattern. Effects related to patients were modest (Bray–Curtis p = 0. 016; Spearman p = 0. 074) (STable 28), while stage effects were more substantial and consistent across measures (Bray–Curtis stratified p = 0. 0024; Spearman stratified p = 0. 0042) (STable 28). Corollary analyses supported these findings, with patient identity accounting for the majority of expression variance (R² ≈ 0.2–0.87; p < 0.001) (SFig. 5C; STable 29), while stage and age had minimal contributions (p > 0.1) (SFig. 5C; STable 29). Composition-based analysis showed a somewhat higher correlation with stage, consistent with overall immune remodeling in metastases. These results indicate that inter-patient heterogeneity predominantly shapes bulk transcriptional profiles. At the same time, the disease stage strongly influences immune composition, especially with increased neutrophil-related programs in metastatic lesions. These data suggest that patient identity primarily influences tumor-intrinsic expression, whereas disease stage more strongly influences immune and stromal composition.

## Materials and Methods

### Dataset assembly for sequencing

Cryopreserved primary appendicular OS and dog-matched metastatic tumor samples were used for whole-genome sequencing (WGS) and single-cell/single-nuclei (sc/sn) RNA sequencing. Samples were collected under approved IACUC protocols as part of ongoing clinical trials at Tufts University Cummings School of Veterinary Medicine. Dog-matched normal tissue (skeletal muscle) was used for germline analysis. Stage-matched cryopreserved plasma samples from 8 dogs were also included.

We assembled a cohort of 22 pet dogs with appendicular OS, comprising 71 samples, including patient-matched primary and metastatic tumors and associated matched cfDNA plasma (Table S1). In the WGS cohort, there were a total of 15 dogs with a primary tumor. The primary tumor from one dog was excluded due to low tumor content and a second dog did not have a primary tumor sample available for analysis. The metastatic samples from these dogs were included in the study, because multiple metastatic lesions were present. Metastatic sites include lung (N=18), subcutaneous (SQ) (N=6), renal (N=3), and secondary bone (N=1). A subset of dogs in the WGS cohort had cfDNA available that was subjected to WGS: cfDNA-1 (primary tumor timepoint; N=8), cfDNA-2 (metastatic timepoint; N=8). A total of 10 primary and 12 metastatic samples from 10 dogs underwent RNAseq. In the RNAseq cohort, all metastatic samples were from the lungs. We evaluated sc/snRNAseq in 6 dogs. This cohort was composed of 4 primary OS tumors, 7 pulmonary metastases and one SQ metastasis. Most dogs had a primary and matched metastatic sample; two libraries captured only one state each (one primary-only, two metastasis-only) after single-nuclei suspensions did not meet quality criteria for subsequent library preparation.

Details of the dog population and clinical treatment are provided in Table S1. A clinical data associated with a subset of samples are reported by Mason et al.^55^ This cohort allowed us to: 1) analyze single-nucleotide variants (SNVs), copy number variants (CNVs) and structural variants (SVs); 2) evaluate genome-wide mutation burden; 3) compare variant landscapes across primary and multiple metastatic tumor tissue sites and plasma within each dog; and 4) characterize tumor-specific evolution of transcriptomic signatures in early and late stage OS.

### Tissue sample WGS library construction and sequencing

Library preparation and sequencing were performed at the Broad Genomic Platform as described. An aliquot of genomic DNA (350ng in 50μL) is used to input into the DNA fragmentation. Library preparation is performed using a commercially available kit provided by KAPA Biosystems (KAPA Hyper Prep without amplification module, product KK8505) and palindromic forked adapters (purchased from Roche). After sample preparation, libraries are quantified using quantitative PCR (kit purchased from KAPA Biosystems) with probes specific to the adapter ends. Based on qPCR quantification, libraries are normalized to 2.2nM and pooled into 24-plexes. Cluster amplification and sequencing were performed on NovaSeq 6000 instruments using sequencing-by-synthesis kits to produce 151 bp paired-end reads. The target depths for Normal and Tumor samples were set at 30x and 60x, respectively.

### cfDNA sample acquisition and extraction

A first set of plasma samples was collected using STRECK tubes from dogs before amputation, and then a follow-up collection was made after clinical metastasis was observed. Frozen plasma (up to 4mL) was extracted using the Qiagen ccfDNA extraction kit, following manufacturer instructions.

### cfDNA library construction and sequencing

DNA input is normalized to 25-52.5 ng in 50 μL of TE buffer (10mM Tris HCl, 1mM EDTA, pH 8.0) according to picogreen quantification. Library preparation uses commercially available KAPA Biosystems (KAPA HyperPrep Kit with Library Amplification product KK8504) and IDT’s duplex UMI adapters. Library quantification was performed using the Invitrogen Quant-It broad-range dsDNA quantification assay kit (Thermo Scientific Catalog: Q33130) with a 1:200 PicoGreen dilution. Following quantification, each library is normalized to a concentration of 35 ng/µL, using Tris-HCl, 10mM, pH 8.0. The samples are pooled and quantified via qPCR and normalized to the appropriate concentration for sequencing. Cluster amplification of library pools was performed according to the manufacturer’s protocol (Illumina) using Exclusion Amplification cluster chemistry. Then, sequencing was performed on a NovaSeq 6000 sequencer, and each pool was run on one lane using paired 151bp runs.

### sc/snRNAseq sample processing, library preparation and sequencing

sc/snRNA-sequencing was performed on 4 primary and 8 metastatic samples from 5 individual dogs using a commercial droplet-based platform (10X Chromium). Briefly, live cell suspensions were obtained from 1g pieces of tissue immediately after collection. Tissue was digested in DNase I (Sigma DN25) and Collagenase II (Gibco 17101015) using a Miltenyi gentleMACS Dissociator with heaters. Cells were passed through a 70 micron cell strainer before histopaque density gradient centrifugation and red blood cell lysis (Biolegend). Cell viability was determined using AO/PI on a Cellometer K2 instrument. 80% cell viability was required for use in library preparation.

Nuclei were obtained from flash-frozen tumor samples. Per the manufacturer’s instructions, the tissue was mechanically pulverized using a Covaris CP02 instrument. Approximately 50mg of pulverized tumor powder was input into the Chromium Nuclei Isolation Kit (10X Genomics; PN-1000493). Nuclei were counted on a Cellometer K2 and visualized under a microscope to confirm nuclei integrity.

Single-cell or single-nuclei suspensions (2000 cells) were input into the 3-prime NextGEM library preparation according to the manufacturer’s instructions on a Chromium controller (STable 30). The resultant libraries underwent quality control and normalization using a Qubit fluorometer and Agilent tapetation. Libraries were pooled and sequenced on a NextSeq 2000.

### Bulk RNAseq sample processing, library preparation, and sequencing

Bulk RNA sequencing was performed on 11 primary and 15 metastatic OS samples collected from 14 individual dogs. Total RNA was isolated from frozen tumor tissue using the Qiagen RNeasy Mini Plus Kit (Qiagen, Hilden, Germany) following the manufacturer’s protocol. RNA quality and integrity were assessed using standard electrophoretic and spectrophotometric methods before library preparation. Sequencing libraries were prepared using the Illumina TruSeq Stranded mRNA Library Preparation Kit (Illumina, San Diego, CA, USA). Libraries were sequenced as 75 bp paired-end reads on an Illumina NextSeq 2000 platform to a target depth of approximately 50 million reads per sample.

### Whole Genome Sequencing Analysis

#### Preprocessing of WGS data

Tumor samples achieved a mean sequencing depth of 73x (range 43-120x). Pre-processing of sequencing data was completed as per Megquier et al., 2022 ^20^. Fastq files were aligned to the canine reference genome (CanFam4 + ROSY) using BWA and underwent quality control in accordance with GATK best practices ^56–62^. For all GATK tools, version 4.2.3.0 was used unless otherwise stated. Duplicate reads were identified using Picard Tools MarkDuplicates (http://broadinstitute.github.io/picard). Base Quality Score Recalibration (BQSR) used a VCF file containing germline variants from 1987 pet dog samples ^61,62^. Tumor fraction was computationally estimated in all 60 samples using the ichorCNA tool ^19,63^.

#### Simple somatic mutation calling

Simple somatic mutations (single-nucleotide variants (SNVs) and small insertions/deletions (indels)) were detected using a robust consensus-calling approach. This method combined three variant callers: Mutect2, Strelka2, and VarScan2 ^64–66^. A VCF containing germline variants was used in the CalculateContamination step. Mutect2 was run with the additional arguments “—downsampling-stride 20—max-reads-per-alignment-start 6—max-suspicious-reads-per-alignment-start 6.” FilterMutectCalls was run with the “—run_orientation_bias_mixture_model_filter” option set to “True” and the “—min-median-read-position” option set to 10 bp. The default settings were used for Strelka2 and VarScan2, but both were run in the tumor-normal configuration. The completed VCF files were processed with bcftools isec to retain only variant calls identified by more than two variant callers, yielding a consensus. We only kept calls that passed filtering using the FILTER flag (FILTER = PASS). These consensus calls were used for all analyses.

#### Structural variant calling

Structural variants (SVs) were called using Manta version 1.6.0. in the tumor-normal configuration^67^. The output VCFs were processed using the Manta-provided script “convertInversion.py” to convert inversions to the older INV format rather than the current break-end (BND) format. The calls were then processed using bcftools, removing any calls that were not passing based on the “FILTER” flag (FILTER != “PASS”) or calls that were marked as imprecise (IMPRECISE = 1). Finally, structural variant calls were converted using the manta-provide script convertInversions.py to convert inversions to the older INV format rather than the current break-end format (BND).

#### Copy number variant calling

Somatic copy number aberrations (SCNAs) were detected using the GATK somatic CNV Pipeline^56,68^. An autosomal panel of normals was created using germline samples from dogs included in this study, and additional male-only and female-only panels were designed for dogs with osteosarcoma. We used the software in tumor-normal mode. We called the autosomes and the X chromosome separately, as recommended by GATK best practices. Copy number losses with a log2 fold change of ≥0.4 (one copy gain) or ≤-0.9 (two copy loss) were considered for analysis. Focal copy number variants were defined as events <3 Mb. For plotting copy number variants, we used PlotModeledSegments in the default settings, plotting autosomes and chromosomes separately and then combining them. The plots containing the raw copy number variant for all samples were included in the supplementary materials. To ensure accurate statistical comparisons across metastatic sites, secondary bone metastasis was excluded as only one sample was present.

#### Oncoprint Visualization

The filtered structural, copy number, and single-nucleotide variants were integrated for visualization using ggplot and cowplot^69,70^. Furthermore, we created an oncoprint using the OncoPrint function in the ComplexHeatmap package, with additional manual modifications in Adobe Illustrator 2023^71,72^. In the oncoprint, we determined the mutations in at least 33% of the samples. Then, we subset this based on the top 20 mutated genes, COSMIC Tier 1 Consensus genes, and recurrent pathways in OS^73,74^. Then, we combined all outputs into a single list of 54 genes. To focus our analysis, we limited single-nucleotide variants to coding regions and restricted them to genes previously identified in canine and human OS. Additionally, we include prevalent genes within the dataset to show newly identified genes of interest in both primary and metastatic samples.

### Chromothripsis Detection

Osteosarcoma is characterized by significant chromothripsis, and we sought to identify these events within our whole-genome sequencing dataset. Existing chromothripsis detection tools, such as ShatterSeek, were developed and validated exclusively on human cancer genomes and have not been tested on canine data. We therefore developed a custom pipeline that implements the criteria described by Korbel & Campbell^75^. We identified events using criteria based on structural variant and copy number variant data from whole-genome sequencing results. Structural variant clusters were identified within 10 Mb windows, requiring ≥10 SVs for consideration of chromothripsis. Copy number oscillation was assessed by counting state transitions within regions containing ≤3 Copy number states, with ≥7 oscillations required for the hallmark pattern. Fragment join randomness was evaluated using a chi-square test against the uniform distribution of the four join types (deletion-like, duplication-like, head-to-head, tail-to-tail). Events were scored based on SV count (1-3 points), CN oscillation (1-3 points), random joins (2 points), and SV type diversity (1 point), with High confidence assigned at ≥7 points and Medium at 5-6 points. Chromosome enrichment was assessed using Poisson tests comparing observed event counts to expected counts based on chromosome length (GCF_011100685.1), with Benjamini-Hochberg correction for multiple testing.

#### WGS Recurrent Pathway Analysis

For general pathway enrichment analysis, we used the STRINGdb to determine enrichment across all pathways. We set the species to dog (species = 9615) and required the score to be over 700^76^. Furthermore, for SVs and CNVs, we needed at least 4 samples or ran the analysis across all genes within a specific stage. We also focused on recurrent pathways in osteosarcoma and performed a general pathway enrichment analysis. Copy number variants (CNVs) were assigned to pathway axes by mapping affected genes to six curated signaling and regulatory groups: PI3K, MAPK, RTK–RAS, NOTCH, TP53, and Epigenetic regulation. Each CNV was classified as a gain, loss, or neutral event based on the mean log₂ copy-ratio threshold (≥ 0.4 = GAIN, ≤ –0.9 = LOSS). Genes were required to recur in at least k patients within a given stage (Primary or Metastatic) and CNV arm (GAIN or LOSS) before inclusion in downstream analyses to minimize the influence of rare or potentially spurious events.

We systematically evaluated k values from 1 to 5 to identify a threshold that balanced recurrence stringency with sample retention. For each k, we recomputed the per-patient paired CNV matrices and measured (i) the number of pathway axes retained (defined as those with ≥ 5 paired patients), (ii) the median paired effect size (mean difference in Metastatic-only versus Primary-only gene counts), (iii) a stability-adjusted score combining effect size and bootstrap confidence-interval width, and (iv) the number of pathways with significant permutation support (label-swap test p < 0.10).

The analysis revealed an elbow at *k = 3*, which preserved the maximum number of supported pathways (six, with permutation *p* < 0.10) while maintaining at least 8 paired individuals per axis. Increasing *k* to 4 or 5 reduced the number of analyzable pairs and weakened significance across most pathways, whereas *k* = 1–2 allowed low-recurrence, potentially spurious events. We therefore selected *k = 3* as the best balance between recurrence and statistical power.

We defined Metastatic-only, Primary-only, and Shared gene sets for each patient and pathway axis, representing genes gained only in the metastatic sample, the primary tumor, or both, respectively. To account for differences in the total number of genes per pathway, we computed a *net-rate* value for each patient, defined as the difference between the number of metastatic-only and primary-only genes divided by the total number of unique genes affected in either or both samples. This metric offers a normalized measure of the relative burden of pathway-specific CNV gains in the metastatic lesion compared to its matched primary.

Paired Wilcoxon signed-rank and *t*-tests assessed whether metastatic lesions had significantly more CNV gains than their matched primaries. The net-rate values were tested against zero using a one-sample Wilcoxon test to see if the median normalized difference was nonzero across patients. Directionality was further evaluated with a two-sided binomial sign test comparing the fraction of patients with greater metastatic-specific CNV burden to the null expectation of 0.5. All *p*-values were corrected for multiple testing using the Benjamini–Hochberg false discovery rate (FDR) procedure, performed separately for gain and loss arms.

#### Distance metrics and PERMANOVA

The similarity between samples in whole-genome sequencing data was evaluated based on gene mutation profiles, focusing on the presence or absence of mutations. A Jaccard index measured the similarity of these profiles across different analyses: 1. Including all samples, 2. Excluding cfDNA samples, and 3. Excluding both cfDNA and renal samples. These analyses aimed to determine whether cfDNA samples contributed to observed dissimilarities, given that they typically contain only a subset of tissue mutations. The limited number of three renal samples could also influence the similarity results. Pairwise similarity matrices were summarized as (i) within-patient versus between-patient comparisons and (ii) within-stage versus between-stage comparisons. These were visualized using violin plots with median lines. To quantify clustering strength, a Δ statistic (the difference between median within-group and between-group similarities) was used, with bootstrap 95% confidence intervals (B = 5,000) computed for the median Δ. Significance was tested with permutation tests (B = 5,000): global permutations for patient effects, stratified within-patient to isolate stage effects, and unstratified for overall stage-related variation. PERMANOVA (adonis2) with Bray–Curtis distances validated the results, reporting marginal R² for patient and stage effects, with blocked permutations by patient for stage-specific tests.

### Single Cell Analysis

#### Single Cell Preprocessing and Filtering

We used the nf-core scrnaseq pipeline v.3.0.0 to process the raw single-cell RNA-seq data^77^. First, raw sequencing files were converted to fastq and demultiplexed using cellranger mkfastq (v.8.0.0). Counts for the expression libraries were derived by using the canine genome (CanFam4 + CanFam3 MT included) ^59,62,78^ using Cellranger count (v.8.0.0) ^79^.

We used an in-house single-cell processing and analysis pipeline to perform quality control, dimensionality reduction, batch correction, clustering, and cell-type annotation. First, we performed quality control (QC) steps to detect and remove cells with low-resolution profiles and those contaminated with ambient RNA using CellBender ^80^. We ran CellBender (remove-background) on the raw 10x matrices per-sample with CUDA support, specifying the expected cell count and droplet range, and running with --fpr 0.01 and --epochs 150. Putative doublets were then detected with DoubletFinder for each sample, and only cells classified as singlets were kept for all downstream analyses. Removing doublets before QC prevents QC thresholds from being inflated by high-complexity doublets.

We performed additional quality control to exclude low-quality cells based on transcript complexity and mitochondrial and ribosomal content. Quality filtering was conducted in a data-driven manner based on global filters. Cells were removed if they had abnormally low numbers of detected genes or UMIs, defined as outliers greater than 1.5 median absolute deviations (MADs) below the median, using the isOutlier function from the scater package ^81^. Cells with unusually low transcript complexity (log₁₀[genes/UMI]) or high mitochondrial content (>3 MADs above the median) were also excluded. Additionally, we capped ribosomal RNA content at the 90th percentile of the per-batch distribution to avoid retaining cells with excessive ribosomal transcript representation.

Cells were retained if they exceeded both thresholds and stayed below the ribo cap; we did not apply upper limits on UMIs and detected genes because DoubletFinder was applied to the data. Doublets typically have more UMIs and detected genes than singlets, so the upper limit would likely include high-complexity singlets. This approach maintains legitimate high-complexity singlets while removing low-quality and high-ribo outliers in a library-aware manner.

#### Dimensionality reduction, batch correction, and clustering

Following QC, gene counts were normalized by total expression, multiplied by a scale factor of 10,000, and log-transformed to compare gene expression between sample cells. Highly variable genes were selected using Seurat’s standard workflow. Principal component analysis (PCA) was performed on the scaled expression matrix using the top 2000 highly variable genes and the top 17 significant PCs, which explain the most variability. We applied the Harmony batch correction method^82^ to the PCA embeddings, using ‘flowcell’ as the batch covariate to account for technical effects from library preparation and sequencing runs. The Harmony-corrected low-dimensional representation was then utilized for downstream graph construction and visualization.

Shared nearest neighbor (SNN) graphs were computed on the top 17 of Harmony embeddings. Segments were identified using Louvain or Leiden community detection, as implemented in Seurat, with a resolution adjusted to ensure stable communities. Two-dimensional embeddings were generated using UMAP in the Harmony space to visualize cell clusters.

We first performed annotation at a broader resolution using SingleR^83^, a reference-based approach for cell type annotation to assign cell type identities to each cluster. SingleR compares the gene expression profiles for each cluster/cell against labeled reference datasets using correlation. For this analysis, we used reference datasets including HumanPrimaryCellAtlasData and DatabaseImmuneCellExpressionData from the celldex R package ^83^. We further performed cell type classification at a finer resolution by partitioning the broader cell types into subtypes using graph-based clustering based on cell type signatures from literature^54,84^. These signatures were used to identify subtypes within each primary cell type through gene set enrichment analysis, applying a hypergeometric test via the HypeR R package^84^. To confirm the cell type labels aligned with expected cell-type specific gene expression patterns, the cell types were inspected based on canonical marker genes (e.g., endothelial: *PECAM1*, *VWF*, *CD34*; B cell: *MS4A1*, *CD79A*; T cell: *CD3D*, *CD3E*, *IL7R*; myeloid: *IL13RA1*, *PPARG*, *CD274*; fibroblast: *ACTA2*, *COL3A1*, *PDGFRB/A*; malignant osteoblast programs: *COL1A1*, *POSTN*, *IBSP*, *BGLAP*) and module scoring of the signatures. In addition, we examined gene expression correlations across different cell types as a consistency check to confirm biological consistency among cell type annotations.

#### Differential gene expression analysis

To identify gene markers distinguishing fine-grained cell types, we set cell identities to *celltype* in the Seurat object. Marker genes for each cluster were computed using Seurat’s FindAllMarkers function on the “RNA” assay, restricted to positive markers (only.pos = TRUE), a log₂ fold-change threshold of 0.25, and a minimum detection rate of 10% (min.pct = 0.1). The top 200 marker genes per cluster, ranked by average log₂ fold-change, were extracted for visualization and annotation. The complete marker list was exported for reference. Custom helper functions were used to standardize gene symbols (clean_sym) and to flag olfactory receptor or trace amine–associated receptor genes, which were excluded from downstream summaries to avoid annotation artifacts.

Compartment-specific differential gene expression analysis between Primary and Metastatic osteosarcoma samples was performed using the Seurat (v5) and MAST frameworks in R. Starting from a preprocessed Seurat object, each cell was assigned to one of five compartments derived from its fine-grained cell-type identity: Tumor (malignant osteoblasts), Myeloid (myeloid, osteoclast, or OC-like clusters), Lymphoid(T and B cells), Stroma (fibroblast and endothelial). The sample type variable was encoded as a factor with “Primary” set as the baseline. For each compartment, differential expression was tested only when both conditions were represented by at least 30 cells. Genes were filtered to exclude mitochondrial transcripts and were retained if expressed in ≥50% of cells in the smaller condition group or in ≥25 cells, whichever threshold was higher. Expression values corresponded to log-normalized counts from the Seurat “RNA” assay. Differential expression was computed using FindMarkers with MAST as the statistical test, including z-scored total RNA counts (nCount_RNA_z) and mitochondrial proportion (percent.mt_z) as latent covariates to control for technical variation. The resulting output contained nominal p-values, Benjamini–Hochberg–adjusted false discovery rates (FDR) to account for multiple testing, log₂ fold changes, and detection fractions per condition. Genes with FDR < 0.05 were considered significantly differentially expressed, with positive log₂FC indicating higher expression in metastasis.

#### Single Cell Geneset Enrichment

To assess the pathway and gene set activity at the single cell level of resolution, we used the singleseqgset R package to perform a gene set enrichment analysis^85^. Expression values were pulled from the RNA assay of the Seurat object. For the preprocessing steps, we used the logFC function, which computes the log fold change for each gene in each cell type relative to all other cells. These logFC values were used as the input for the wmw_gsea function to perform the gene set enrichment analysis, which applies a Wilcoxon-Mann-Whitney rank sum test, which evaluates if each gene in a gene set is systematically up- or down-regulated relative to all other genes within a cluster.

For each gene set and cluster, enrichment statistics and p-values were obtained from the wmw_gsea() function. The resulting p-values were subsequently adjusted for multiple hypothesis comparisons using Benjamini–Hochberg false discovery rate (FDR) correction. Gene sets with an FDR-adjusted p-value < 0.05 were considered significantly enriched.

We visualized the results using z-scoring across gene sets (row-scaled) to emphasize differences in pathways across the cell types. Then, heatmaps were generated using ComplexHeatmap package with hierarchical clustering applied to both rows and columns^71^. Additionally, signed bar plots were produced to summarize the top gene sets per cluster.

#### Distance metrics and PERMANOVA

To evaluate similarity between samples in the single-cell data, we used pseudobulk expression profiles and cell-type composition estimates. Pseudobulk profiles were generated by averaging gene expression across samples for the top 2,000 most variable genes, capturing key transcriptional differences and reducing noise from low-variance genes. Cell-type composition was determined from the proportional representation of annotated cell types per sample, providing a complementary view of the microenvironment. For each dataset, we calculated Bray–Curtis similarity (1−DBray) using vegan::vegdist, which measures magnitude-based proportional differences, and Spearman correlation, which assesses rank-based agreement regardless of scale. This dual approach helped us see if sample organization was consistent when focusing on expression magnitude or relative order. These metrics provide different insights, as cell-type proportions are compositional and pseudobulk expression is continuous. We summarized pairwise similarity matrices as (i) within-patient vs. between-patient and (ii) within-stage vs. between-stage groupings, visualized with kernel density plots and median markers. A Δ statistic for each sample, calculated as the difference between within-group and between-group mean similarities, quantifies the strength of clustering. Bootstrap (B=5,000) provided 95% confidence intervals for the median Δ. Significance was tested with permutation tests (B=5,000): global permutations for patient effects, stratified within-patient to isolate stage effects beyond patient identity, and unstratified for overall stage differences. PERMANOVA (adonis2) on Bray–Curtis distances validated these findings, reporting marginal R² for patient and stage effects, with blocked permutations by patient for stage-specific analysis^86^.

### Bulk RNA Sequencing Analysis

#### Bulk RNA sequencing preprocessing

Raw paired-end FASTQ files (75 bp) were processed using the nf-core/rnaseq pipeline ^87^ with the STAR–RSEM workflow. Adapter sequences and low-quality bases were removed using Trim Galore! ^88^, and read quality was assessed before and after trimming with FastQC ^89^. Alignment was performed with STAR (two-pass mode) against the *Canis lupus familiaris* reference genome (UU_Cfam_GSD_1.0_ROSY) ^59,62^, followed by sorting and indexing with samtools. Gene- and transcript-level quantification was performed with RSEM ^90^, which estimated expected counts and transcripts per million (TPM) using the STAR alignments and the corresponding gene annotation ^91^. Quality metrics, including mapping efficiency, duplication rate, and gene body coverage, were compiled and summarized with MultiQC^92^. The final outputs, expected counts, and TPM matrices, were imported into R for normalization, differential expression analysis, and integration with single-cell transcriptomic data.

#### Differential gene expression

RSEM gene-level results were imported with tximport to maintain proper count/length scaling and combined with curated sample metadata. Before modeling, genes with ambiguous identifiers (e.g., LOC-prefixed) were excluded, identifiers were made unique, and lowly expressed genes were filtered using edgeR’s filterByExpr (design-aware). Library sizes were normalized using TMM. We applied a limma-voom framework with quality weights (voomWithQualityWeights) for differential expression analysis. To account for the paired study design, we estimated the within-subject correlation with duplicateCorrelation and included Individual_ID as a blocking factor. Only dogs with both primary and metastatic tumors were included, resulting in 22 samples processed through the DGE pipeline. The design incorporated the biological condition (Metastatic vs Primary) and a technical covariate (Flowcell) to reduce batch effects. After fitting the model (lmFit) and creating the Metastatic − Primary contrast (makeContrasts), empirical Bayes moderation with robust variance estimation (eBayes(robust=TRUE)) was applied. Genes with FDR < 0.05 and |log2FC| ≥ 1 were identified as differentially expressed. The resulting tables (all genes, as well as up- and down-regulated subsets) were exported for downstream visualization and pathway analysis. Visualization was performed using ComplexHeatmap in R, and the results were later mapped onto the single-cell RNA-seq dataset using UCell and visualized with pheatmap.

#### Immune/stromal cell abundance estimation with MCP-Counter

Gene-level RSEM outputs were imported with tximport and converted to counts per sample^90^. After removing ambiguous identifiers (e.g., *LOC*-prefixed genes) and enforcing unique row names, library sizes were normalized with edgeR’s TMM (no batch correction applied at this stage for abundance estimation)^93^. For visualization-only QC, TPM values were also computed, and a light expression filter was applied (TPM ≥ 1 in ≥ 20% of samples). MCPCount calculations used the uncorrected log2-CPM matrix derived from TMM-normalized counts, consistent with the authors’ recommendations for gene-signature scoring in bulk RNA-seq. Gene identifiers were mapped to symbols when needed, and duplicates were collapsed to a single value per gene (median). MCPCounter^94^ was run with featuresType = “HUGO_symbols” and the package’s default marker gene sets to estimate abundance scores for significant immune and stromal populations (e.g., T cells, B lineage, myeloid, fibroblasts, endothelial). The resulting sample × cell-type score matrix was transformed to row z-scores for heatmap display using ComplexHeatmap (row centering and scaling by the row standard deviation; NA and non-finite values were checked and handled conservatively).

### Mixed-effects modeling of MCPCounter cell-type scores

We employed linear mixed-effects models of the form score ∼ condition + (1|patient_id) for each MCPCounter cell type to detect cell populations associated with disease stage, accounting for repeated measures within individuals. In these models, condition denotes the disease stage (Primary or Metastatic), and patient_id was included as a random intercept to account for individual-level variability. This approach remains effective even with unbalanced sampling, such as multiple samples per stage or missing pairs. The model estimates the fixed effect of metastasis compared to primary tumors for each cell type, showing the direction and extent of change (Metastatic > Primary or vice versa). P-values are obtained from the fixed-effect term and adjusted across cell types using the Benjamini–Hochberg method to control the false discovery rate (FDR). After adjusting for within-patient correlation, the coefficients reflect average differences in cell-type abundance across disease stages.

#### Distance metrics and PERMANOVA

Analogous analyses were performed on bulk RNA-seq data using (i) gene expression matrices and (ii) cell-type composition estimated by MCPCounter. The same pair of similarity metrics, Bray–Curtis and Spearman, was used to capture complementary information: Bray–Curtis emphasizing abundance magnitude and Spearman emphasizing rank-based concordance. Because MCP-Counter scores represent semi-quantitative enrichments rather than strict proportions, including both measures helped assess the robustness of the inferred structure. We also assessed the overall shift in similarity distributions using Bray–Curtis and Spearman metrics, which capture different aspects of sample similarity. Bray–Curtis emphasizes proportional differences and is better suited for composition data, while Spearman correlation focuses on relative rankings and is more appropriate for expression data. By pooling similarity scores across both data types, we examined whether overall trends were consistent across metrics and datasets. Similarity matrices were summarized and tested exactly as described for the single-cell data. Visualization used ComplexHeatmap (z-scored pseudobulk or MCP-Counter data, columns split by *Patient_ID* or *Tumor Type*), ggplot2 for violin and density plots, and patchwork/cowplot for multi-panel figure assembly.

#### Bulk RNAseq cell type deconvolution

To estimate the relative abundance of cell populations within bulk RNA-seq samples, we used reference-based deconvolution with MuSiC (v0.2.0)^95^. This method leverages single-cell transcriptomic data to infer cell-type proportions while considering subject-specific variability. The single-cell reference was directly derived from the annotated osteosarcoma dataset used in this study. Gene counts were extracted from Seurat’s default “RNA” assay, and genes lacking valid symbols were removed. Duplicates were collapsed by summing counts to keep one row per gene. Metadata for cell type (celltype) and sample ID (sample_ID) were retained to construct a SingleCellExperiment object, essential for MuSiC. Bulk gene counts were imported via tximport (type=“rsem”, countsFromAbundance=“no”) and rounded to integers. Gene symbols were harmonized to match the reference. Deconvolution involved restricting both datasets to their common gene set and fitting the MuSiC model using MuSiC::music_prop, with parameters clusters=“celltype”, samples=“sample_ID”, and data.type=“RNAseq”. MuSiC employs a weighted non-negative least-squares approach, modeling each bulk sample as a mixture of single-cell profiles grouped by patient and cell type. Genes with lower cross-subject variability are upweighted to enhance stability, and the optimization continues until convergence (iter.max=1000, nu=1e−3, epsilon=1e−3). The resulting cell-type proportions are non-negative and sum to about one per sample, reflecting the relative contributions of each major cell population.

#### Statistical testing of immune differences by stage

We first standardized cell-type labels across the MuSiC output to harmonize related immune subsets and evaluate whether the deconvolved immune compartment differed between primary and metastatic tumors. Columns corresponding to Myeloid, B-cell, and T-cell populations (the latter also encompassing NK and cytotoxic lymphocyte subsets) were identified by regular-expression pattern matching and combined by summing their proportional estimates within each sample. The overall immune proportion was defined as the sum of these three compartments and transformed using a logit link after bounding proportions within [10^−6, 1−10^−6] to ensure numerical stability. For descriptive comparison, values were aggregated at the patient × stage level by taking the median on the logit scale and then back-transforming to the proportion scale for visualization. Differences in immune composition between stages were first assessed using a paired Wilcoxon signed-rank test on matched primary–metastatic pairs, testing whether the immune compartment was higher in metastatic samples. We also fit a linear mixed-effects model to account for repeated measures and patient-specific variation. The logit-transformed immune proportion was modeled as a function of disease stage (Primary vs. Metastatic) with a random intercept for each patient. Specifically, the model estimated immune proportion on the log-odds scale as a function of stage, allowing each patient to have their own baseline immune level. Using lme4 and lmerTest with *condition* (Primary as reference) as a fixed effect and a random intercept for the patient, estimated marginal means and contrasts were obtained using emmeans, with back-transformation to the probability scale for interpretation. We reported odds ratios (ORs) and 95% confidence intervals derived from these contrasts, which were exponentiated from the logit-scale estimates. These complementary non-parametric and mixed-effects methods provided robust inference on stage-related shifts in the deconvolved immune composition while accounting for within-patient correlation.

## Discussion

While multiple studies have charted the genomic and transcriptomic features of primary and metastatic OS^10,14,18,96^, the pervasive genomic heterogeneity across patients underscores the need for longitudinal assessment of genetic features to identify collateral sensitivities within patients. By integrating pet dog matched metastatic and primary OS samples across multiple sites and three sequencing modalities (WGS, bulk RNAseq, scRNAseq) in the present study, we resolved patient-dependent tumor evolution and microenvironment remodeling across all stages of disease at both individual and population levels. Our results demonstrate that differences between patients are driven by shared clonal ancestry and tumor-specific pathways and outweigh spatial or temporal variability within individuals. While metastatic samples accumulate more mutations than primary tumors, overall mutational composition remains dominated by patient identity rather than anatomic site or disease stage. Therefore, although cross-sectional studies offer valuable insights into overall mutational landscapes, patient-matched and longitudinal designs enhance the ability to identify mutations that emerge over time and differentiate between effects specific to individual patients and those related to disease stage.

Mutation burden increased with disease progression in our dataset. Metastases, particularly lung metastases, exhibited higher tumor mutation burden than primary tumors, although consistent with reports in human OS^14^, we did not observe changes in CNVs or SVs. However, we did identify increased genomic instability in lung metastases compared with primary tumors, which may reflect early chromothripsis events followed by a steady accumulation of mutations as the disease evolves and progresses, suggesting a stage-linked rise in chromosomal instability. Together with a high frequency of DNA damage response (DDR) pathway mutations (e.g., *ATR*, *ATM*, *BRCA1/2*, *PALB2*), these findings implicate replication stress and faulty repair as a pervasive component in the metastatic process. Additionally, the recurrence of DDR pathway dysregulation and chromothripsis in advanced stages of pet dog OS is consistent data from OS in people^10^.

The increased mutation burden in metastatic tumor tissue was recapitulated in cfDNA, using time points collected prior to amputation and following clinical evidence of metastasis. Our data demonstrate a substantial shift in mutations that are more similar to those present in metastatic tissues when compared to those identified in primary tumors, a pattern consistent across both CNV and SNVs. These findings align with previous work in other cancer types demonstrating that serial cfDNA sampling can detect unique mutations in metastatic tumors that were absent in the primary tumor ^97–101^. This supports cfDNA as a minimally invasive tool to monitor disease progression and emerging therapeutic resistance without repeated biopsies.

Across all sequencing modalities, we found evidence of strong inter-patient genomic tumor heterogeneity, underscoring the importance of evaluating OS tumors in a patient-specific context. Using scRNA-seq, we identified fine-scale differences across samples, stages, and patients. Specifically, we observed heterogeneity in the six malignant osteoblast subtypes, where different programs were clearly identifiable, such as proliferative (MOB3/5/6), immune and stress response (MOB1/4), and osteoclast-like and bone remodeling (MOB2). Furthermore, these malignant osteoblasts were present across both primary and metastatic OS stages and patient identity dominated each MOB subtype.

Immune composition exhibited more phenotypic variability across bulk and scRNA-seq datasets. Deconvolution of bulk RNA-seq revealed stage-associated differences in immune composition, with significantly higher T, B, neutrophil and other myeloid cell populations in metastatic samples. This signal was not observed in the scRNA-seq data. The reason for this is likely multifactorial, including differences in dataset processing, small sample size resulting in type II errors and, differences in medical management of dogs in the study that include various immunotherapies impacting myeloid cells across each dataset. Interestingly, neutrophils have been implicated as playing an important role in seeding new metastatic sites/niches as well as promoting local immunosuppression in OS ^102–104^. In support of this, previous studies have demonstrated increased numbers of immune cells, including T cells, B cells, and macrophages, in pet dog and human pulmonary metastases ^105,106^.

While integrating multiple sequencing modalities provided insights into the tumor microenvironment and tumor evolution, we did not perform WGS, bulk RNA-seq, and scRNA-seq on all matched tissue pairs, which limited cross-modality comparisons for individual patients. Furthermore, we could not be confident in structural variants called in cfDNA due to the inherent fragmentation of this sample type. Future work may include other sequencing modalities, such as methylation and ATAC-seq, given that the genes mutated are often involved in epigenetic regulation and could provide novel targets for therapeutic intervention.

## Conclusions

These data provide a framework for the tumor microenvironment and tumor evolution across samples from the same patient. We demonstrate that (i) patient-specific OS tumor evolution governs the tumor-intrinsic genome and transcriptome; (ii) metastatic progression shows increased genomic instability and convergent pathway amplification, and (iii) stage drives the microenvironment remodeling, particularly inflammatory and neutrophil-associated programs. The comparative framework used in this study underscores the importance of cfDNA-tissue profiling for longitudinal monitoring of tumor progression and identification of pathway-level recurrent alterations that can be leveraged for therapeutic intervention.

## Data Availability

The sequence read data in the cram and fastq format for all samples are available in the NCBI Sequence Read Archive (SRA) under BioProject PRJNA1377626.

## Acknowledgements

The authors thank the Broad Genomics Platform and Tufts Comparative Pathology and Genomics Shared Resource for their assistance with library preparation and sequencing. This work was supported by the following grants: ACS Research Scholar Grant, American Kennel Club Companion Health Fund (02768), Morris Animal Foundation (D21CA-026), NIH CTSA UM1TR004398, NIH U01CA224182, Intramural Program of the NCI, NIH (Z01-BC006161), NCI R37CA244355. HLG was supported by K01OD028268-01A1; CP was supported by K01OD34451. DSS was supported by K08 5K08CA290061-02. KM was supported by K01OD036469-01A1. The content is solely the responsibility of the authors and does not necessarily represent the official views of the National Institutes of Health.

## Author contributions statement

CH (data analysis, manuscript writing, review and editing); KS (data generation, manuscript review and editing); TTK (data analysis support, manuscript review and editing); CP (histologic data analysis, manuscript review and editing); KM (data analysis support, manuscript review and editing); DG (manuscript review and editing); JR (data analysis support; manuscript review and editing); EK (manuscript review and editing); DSS (manuscript review and editing); BC (project conceptualization; manuscript review and editing); CAL (provided biosamples; manuscript review and editing); HLG (project conceptualization; data analysis; manuscript review and editing).

---

**Supplementary Figure 1.**
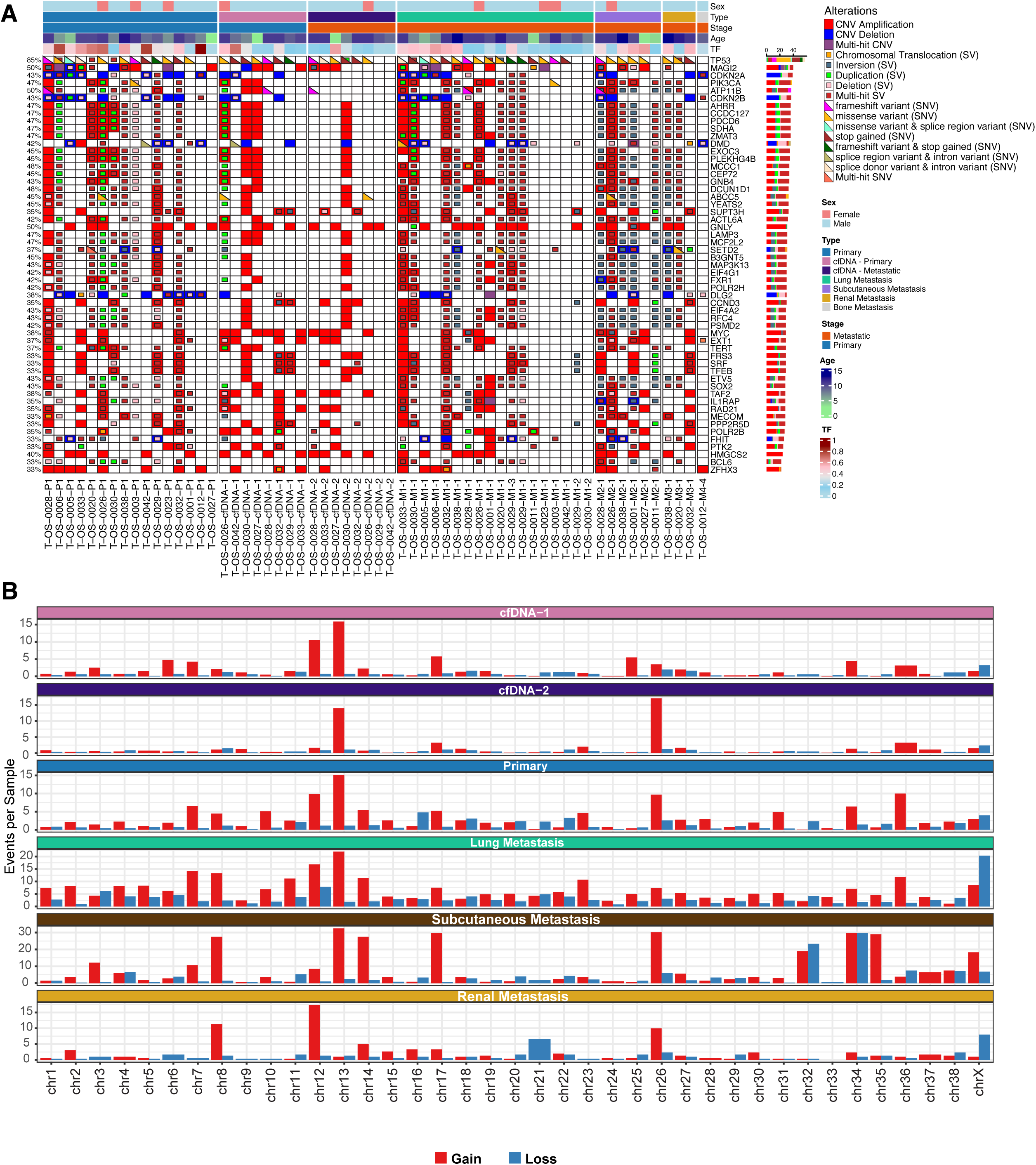
Genomic landscape and mutational burden across cfDNA, primary, and metastatic osteosarcoma samples, clustered by metastatic site. **(A)** Oncoprint showing mutations present in at least 33% of samples, subsetted to the top 20 mutated genes, COSMIC Tier 1 Consensus genes, and recurrent osteosarcoma-associated pathways. The heatmap is clustered by anatomical site (Primary, Lung Metastasis, Renal Metastasis, and Subcutaneous Metastasis) and includes single-nucleotide variants (SNVs), copy-number variants (CNVs), and structural variants (SVs). Top annotations indicate sex, sample type, age, and tumor fraction (TF). **(B)** Copy number events per sample for each chromosome, stratified by sample type. The number of gain (red) and loss (blue) events per chromosome was normalized by the number of samples in each group. Sample types include circulating free DNA at initial (cfDNA-1) and subsequent (cfDNA-2) collection timepoints, primary tumors, and metastases from lung, subcutaneous, and renal sites. Note that y-axis scales differ between panels. Recurrent gains on chromosome 13 are observed across all sample types. Chromosome X losses are particularly prominent in lung metastases.

**Supplementary Figure 2.**
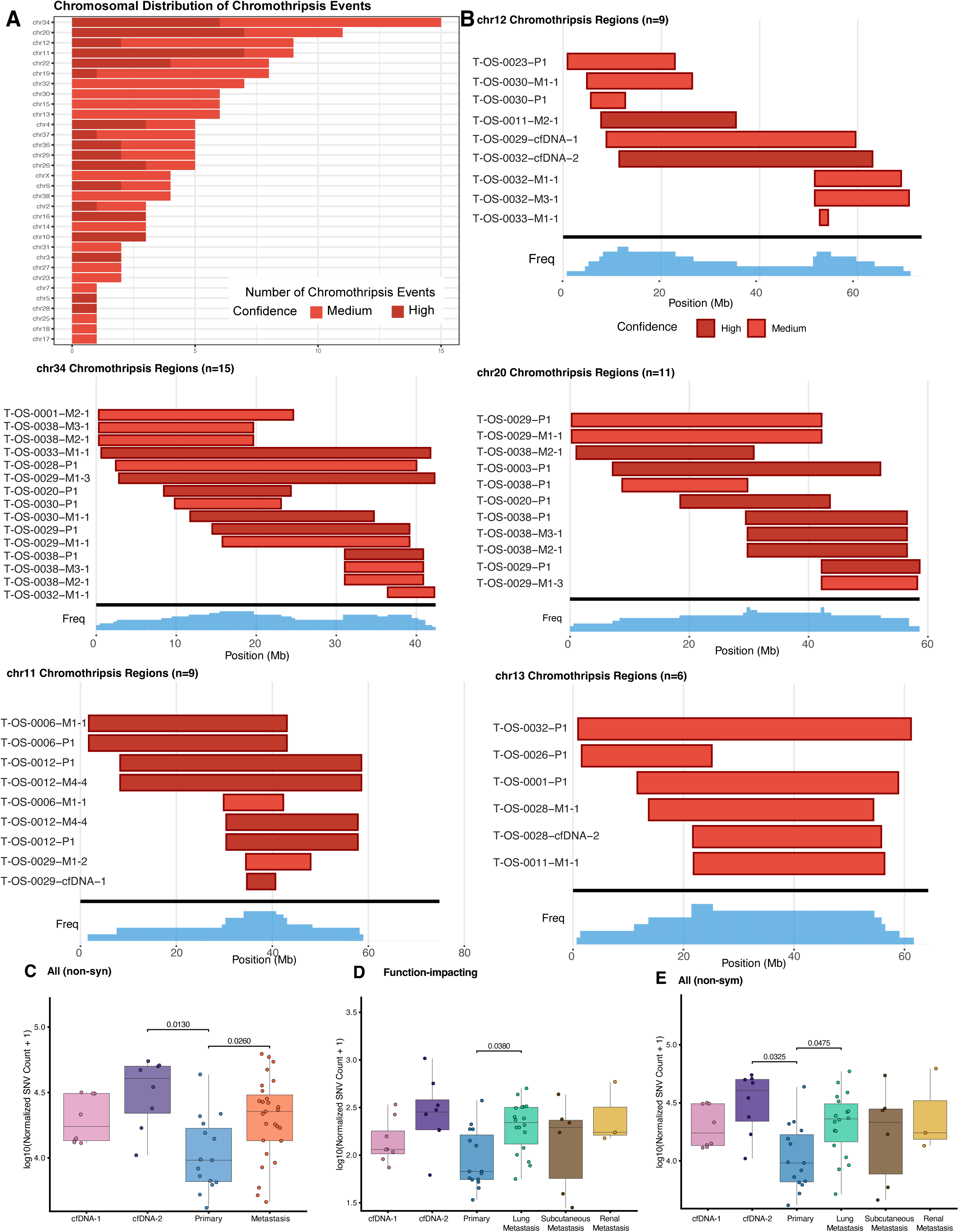
**Chromothripsis patterns and SNV burden across sample types in canine osteosarcoma**. **(A)** Chromosomal distribution of chromothripsis events across all samples. Bar color indicates confidence level of chromothripsis calls (high confidence, dark red; medium confidence, light red). Chromosomes 34, 12, 20, 11, and 13 harbored the most frequent chromothripsis events. **(B)** Genomic coordinates of chromothripsis regions on the five most frequently affected chromosomes. Each horizontal bar represents a chromothripsis event in an individual sample, with bar color indicating confidence level. Sample identifiers denote tumor type (P1, primary; M, metastasis; cfDNA, circulating free DNA). Frequency histograms below each panel show the distribution of chromothripsis breakpoints across the chromosome. The number of chromothripsis events per chromosome is indicated in panel titles. **(C)** Normalized non-synonymous SNV burden compared between cfDNA and tissue samples. Metastases showed significantly higher SNV counts than primary tumors (p=0.0260), and cfDNA-2 samples showed significantly elevated counts compared to primaries (p=0.0130). **(D)** Normalized function-impacting SNV burden stratified by sample type. Lung metastases exhibited significantly higher coding mutation burden compared to primary tumors (p=0.0380). **(E)** Normalized non-synonymous SNV burden stratified by metastatic site. cfDNA-2 samples showed significantly higher SNV counts compared to primary tumors (p=0.0475), and lung metastases showed significantly elevated counts compared to primaries (p=0.0325).

**Supplementary Figure 3.**
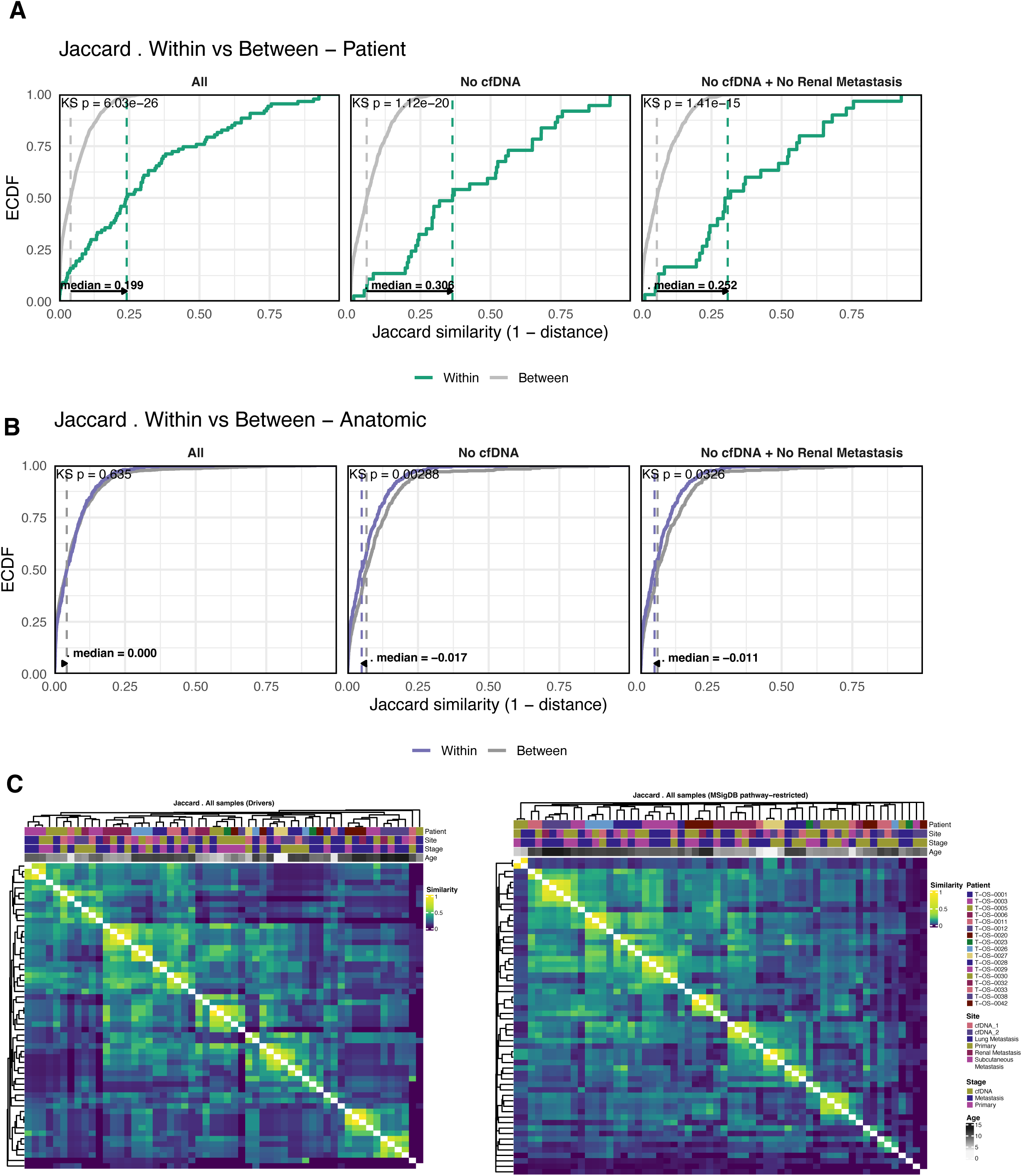
Within- and between-sample mutational similarity and clustering based on whole-genome sequencing data. **(A)** Empirical cumulative distribution functions (ECDFs) comparing Jaccard similarity of shared mutations within and between patients across three sample subsets: all samples (left), excluding cfDNA (middle), and excluding both cfDNA and renal metastases (right). Within-patient comparisons showed consistently higher similarity than between-patient comparisons across all subsets (Kolmogorov–Smirnov p=6.03×10⁻²⁶, p=1.12×10⁻²⁰, and p=1.41×10⁻¹⁵, respectively). **(B)** ECDFs of Jaccard similarity comparing within- versus between-anatomic site comparisons using the same subset groupings. When all samples were included, no significant difference was observed (p=0.635, median difference=0.000). After excluding cfDNA samples, between-site comparisons showed slightly higher similarity than within-site comparisons (p=0.00288, median difference=-0.017), a pattern that persisted when renal metastases were also excluded (p=0.0326, median difference=-0.011). This indicates that mutation profiles are not enriched for site-specific patterns and that patient identity is the primary driver of mutational similarity. **(C)** Pairwise Jaccard similarity heatmaps across all WGS samples using putative driver mutations (left) and MSigDB pathway-restricted variants (right). Samples cluster predominantly by patient rather than anatomic site or disease stage, indicating that mutation profiles are patient-specific and stable across disease contexts.

**Supplementary Figure 4.**
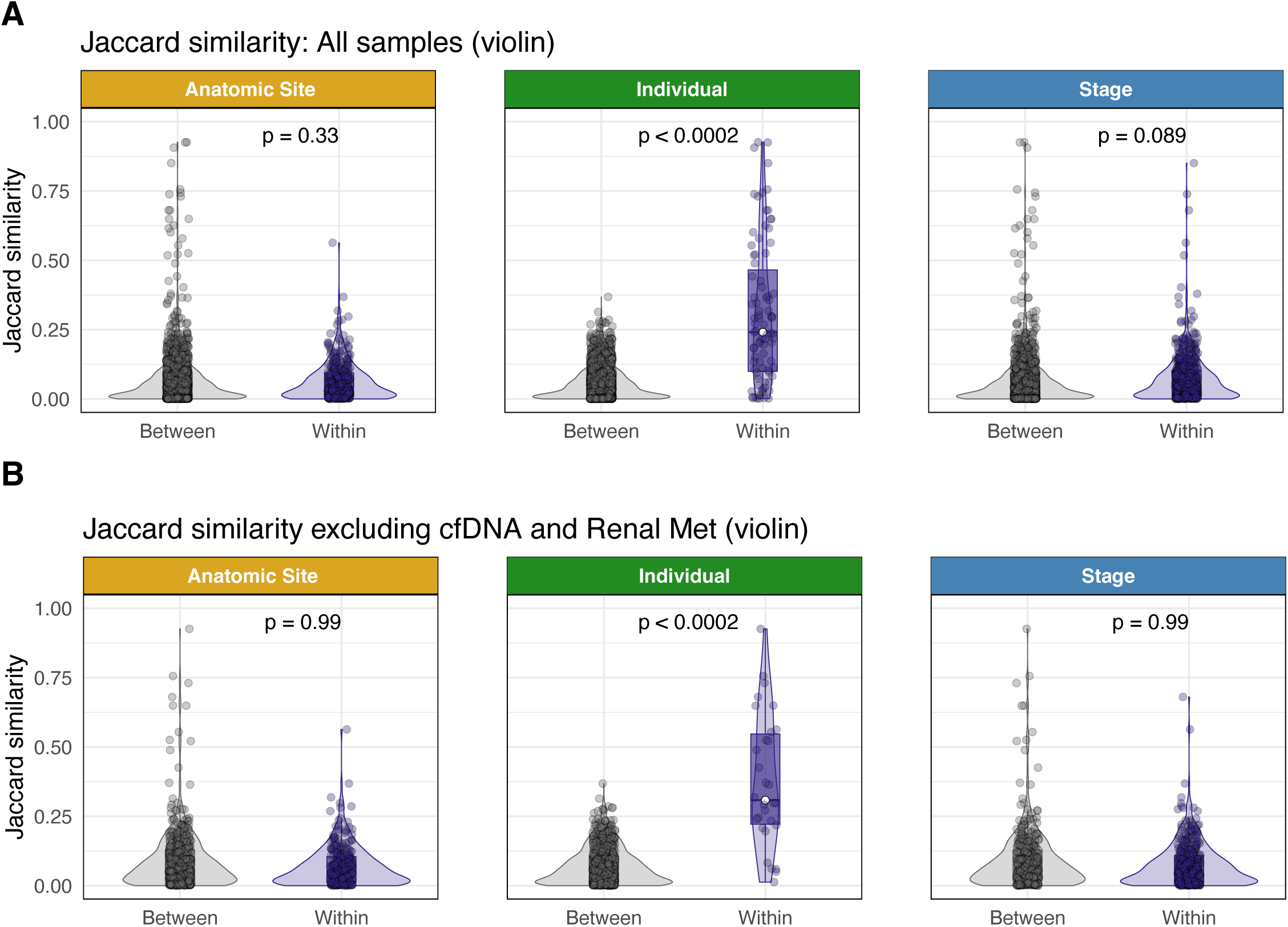
Jaccard similarity of shared mutations across anatomical site, individual, and disease stage. **(A)** Violin plots showing Jaccard similarity scores of shared mutations within and between groups across anatomical site (left), individual (center), and stage (right). Samples from the same individual exhibited significantly higher similarity compared to those from different individuals (permutation, p < 0.0002), while no significant differences were observed for anatomical site (p = 0.33) or stage (*p* = 0.089). **(B)** Jaccard similarity analysis excluding cfDNA and renal metastasis samples. The strong within-individual similarity remained significant (p < 0.0002), whereas site- and stage-based comparisons showed no detectable differences (p = 0.99 for both), indicating that interindividual variation is the primary driver of mutational similarity patterns.

**Supplementary Figure 5.**
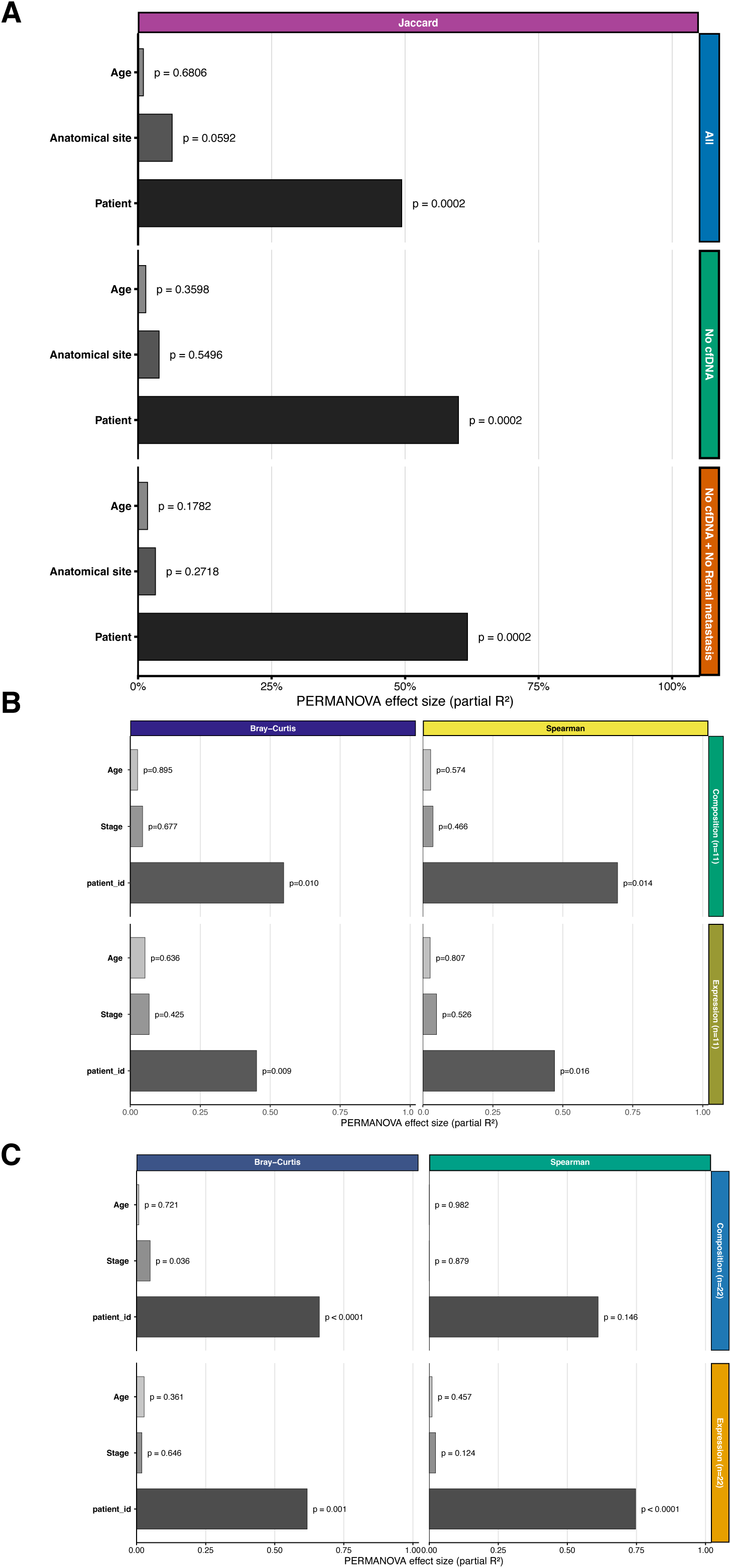
PERMANOVA analysis of mutational similarity and transcriptional/compositional variation. **(A)** PERMANOVA partial *R²* effect sizes based on Jaccard similarity of shared variants, shown for all samples, excluding cfDNA, and excluding cfDNA plus renal metastases. Patient identity accounts for the largest share of variance (see panel labels for *p*-values). **(B)** Single-cell analyses: PERMANOVA using Bray–Curtis (left) and Spearman (right) on pseudobulk scRNA-seq expression (top) and cell-type composition from scRNA-seq (bottom). Patient identity is the dominant factor; stage and age contribute minimally. **(C)** Bulk RNA-seq analyses: PERMANOVA using Bray–Curtis (left) and Spearman (right) on bulk expression (top) and MCPCounter composition (bottom). Patient identity explains the greatest variance across all comparisons. Stage showed a modest significant association with Bray–Curtis expression (p=0.038), but effects of stage and age were otherwise small or non-significant(see panel labels for additional *p*-values).

**Supplementary Figure 6.**
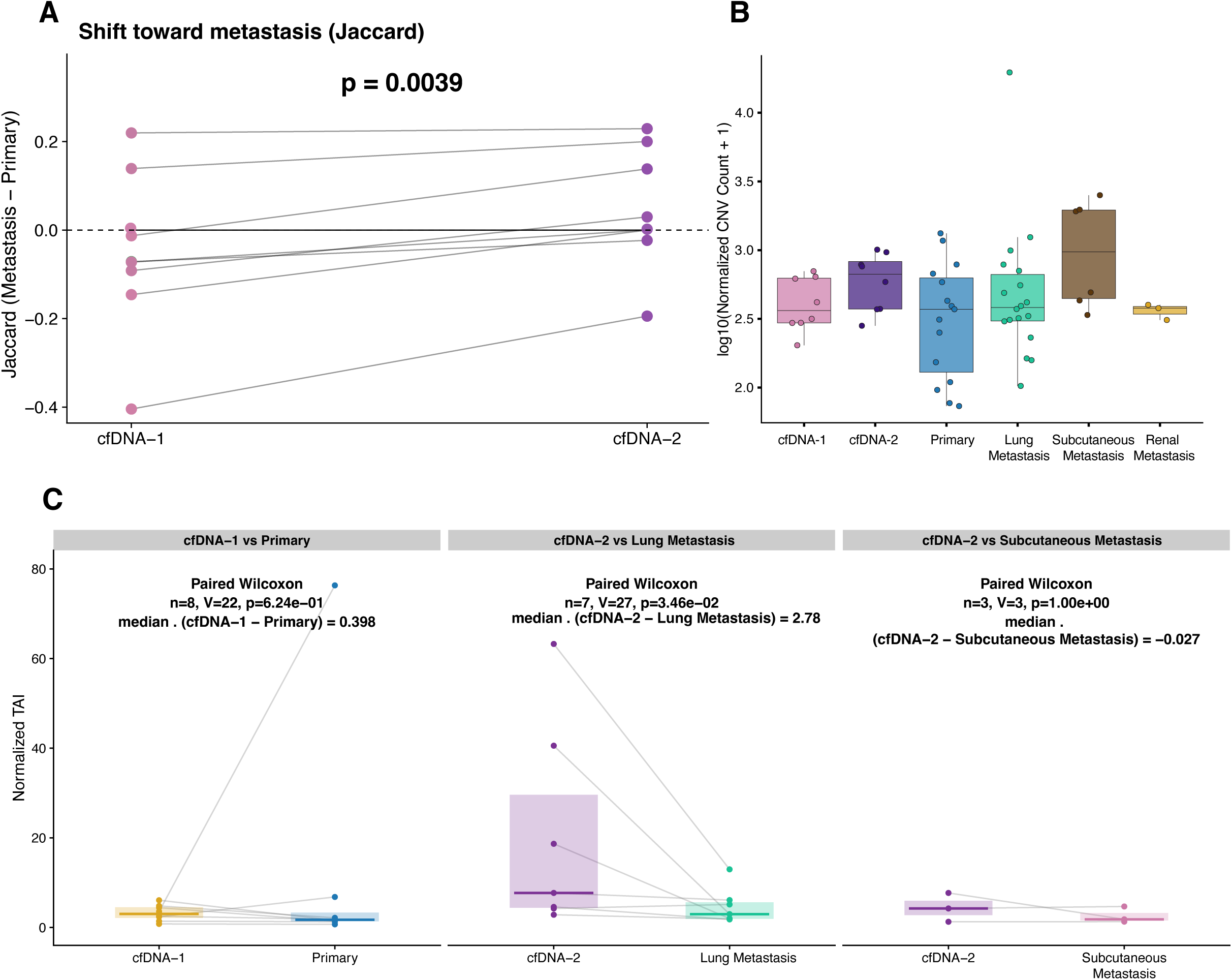
Somatic mutation shift and genomic instability across sample types and metastatic sites. **(A)** Jaccard similarity analysis comparing cfDNA-1 and cfDNA-2 relative to primary versus metastatic mutations. A significant shift toward greater similarity with metastatic profiles was observed in cfDNA-2 compared to cfDNA-1 (p=0.0039), indicating progressive accumulation of metastasis-associated mutations over time. **(B)** Boxplots showing copy-number variant (CNV) counts normalized by tumor fraction across sample types. CNV burden did not differ significantly between groups but exhibited site-specific variability, with lung metastases showing higher median counts. **(C)** Paired comparisons of genomic instability (TAI) between cfDNA and matched tumor samples. cfDNA-1 did not differ significantly from primary tumors (p=0.62). cfDNA-2 exhibited significantly higher TAI relative to lung metastases (p=0.035) but not subcutaneous metastases (p=1.0).

**Supplementary Figure 7.**
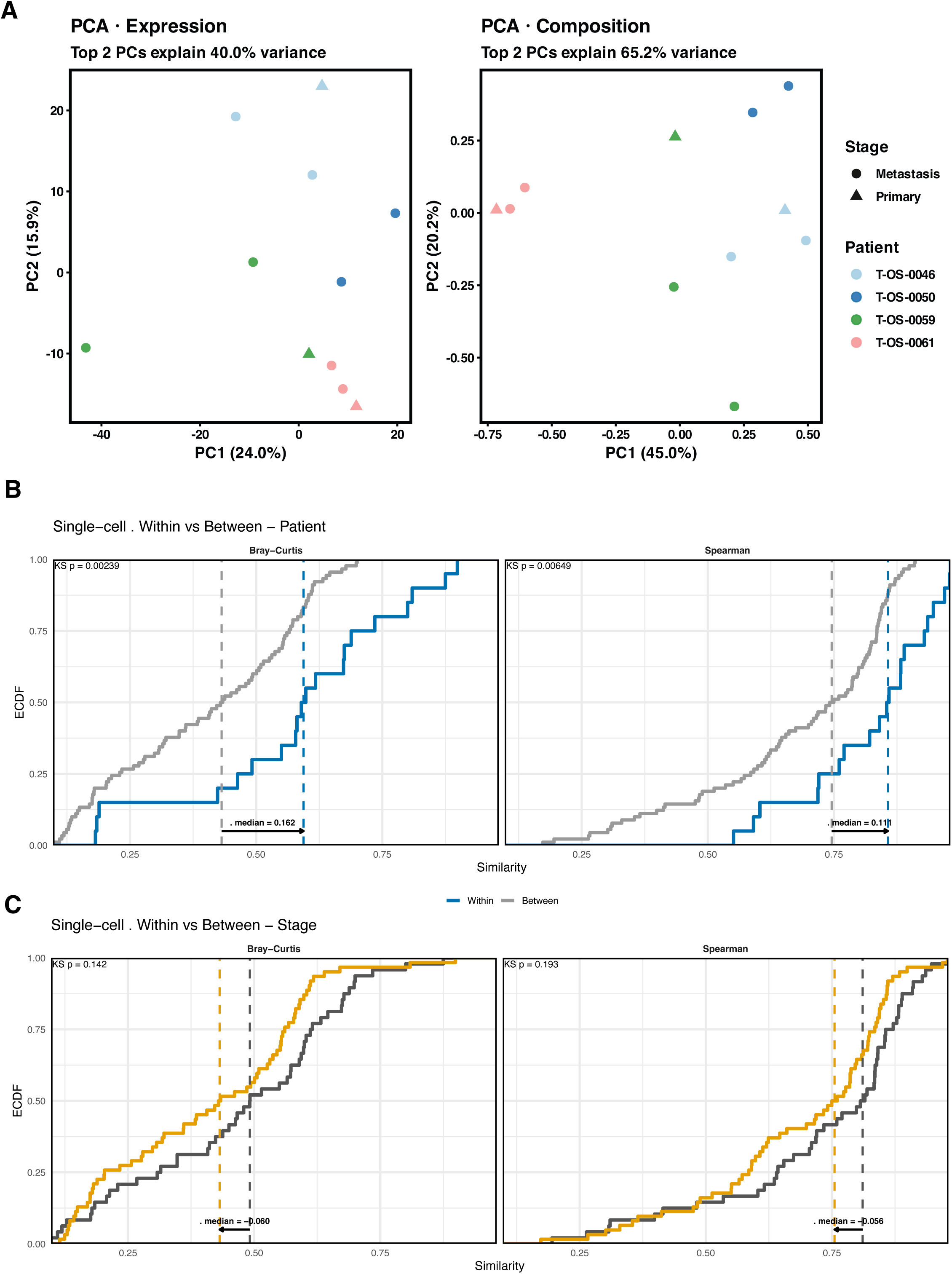
Similarity and principal component analysis of single-cell expression and composition data. **(A)** Principal component analysis (PCA) plots of pseudobulk expression (left) and Hellinger-transformed cell-type composition (right). Samples cluster primarily by patient rather than stage, with the first two principal components explaining 40.0% of the variance for expression and 65.2% for composition. **(B)** Empirical cumulative distribution functions (ECDFs) comparing within- and between-patient similarity in single-cell pseudobulk expression (Bray–Curtis, left) and cell-type composition (Spearman, right). Samples from the same patient exhibited significantly higher similarity than those from different patients (Bray–Curtis *p* = 0.00239; Spearman *p* = 0.00649), indicating strong patient-specific transcriptional and compositional patterns. **(C)** ECDFs comparing within- and between-stage similarity (primary vs. metastatic) for the same datasets. No significant stage-dependent differences were observed (Bray–Curtis *p* = 0.142; Spearman *p* = 0.193), suggesting that interpatient variation outweighs stage-specific effects at the single-cell level.

**Supplementary Figure 8.**
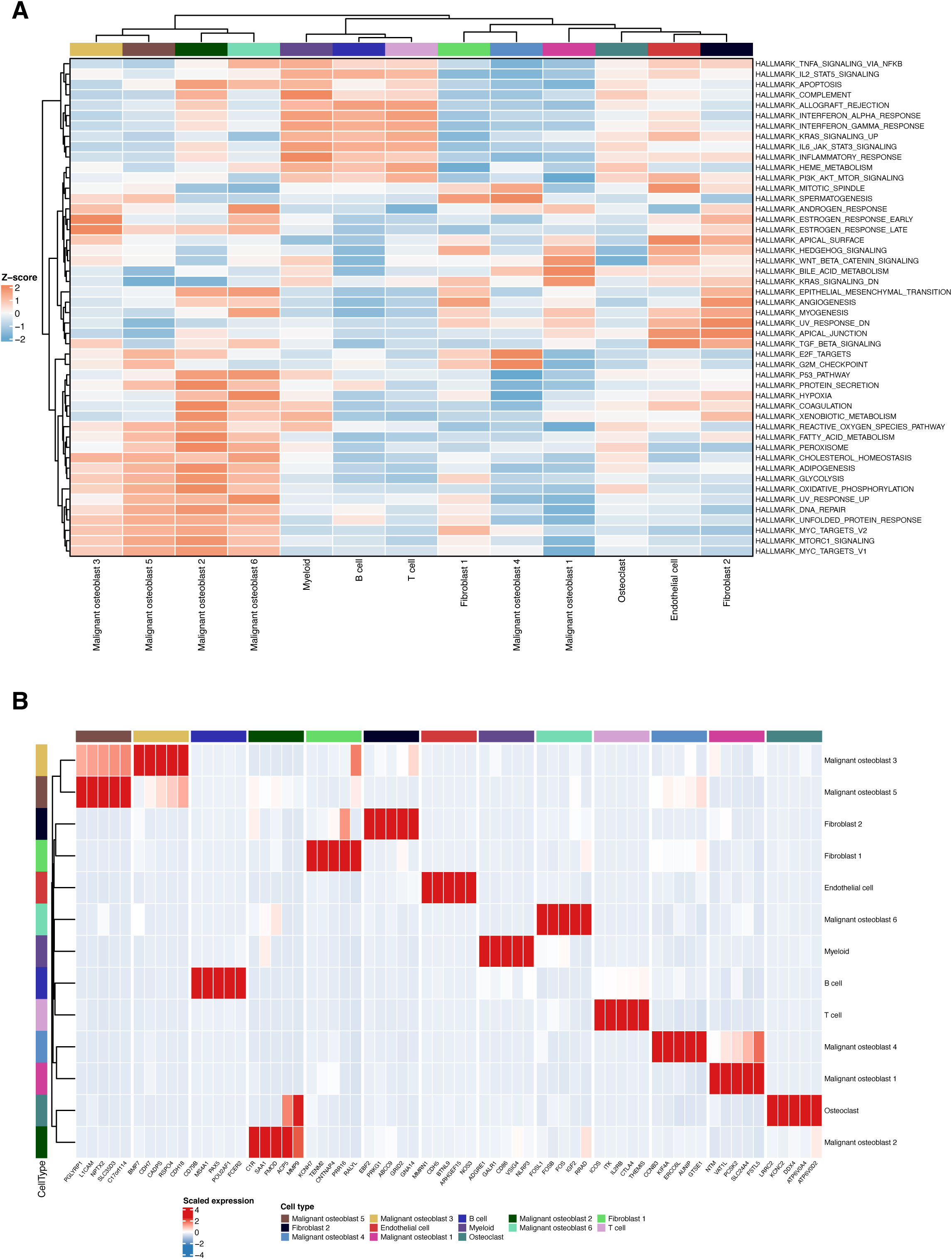
Pathway enrichment and transcriptional signatures across single-cell–defined osteosarcoma cell types. **(A)** Hierarchical clustering heatmap of Hallmark pathway enrichment scores across annotated single-cell populations. Rows represent Hallmark gene sets, and columns represent individual cell types, including malignant osteoblast subtypes (1–6), myeloid, lymphoid, fibroblast, osteoclast, and endothelial populations. Enrichment scores are Z-score–scaled across cell types. Distinct pathway activation patterns are observed among malignant osteoblast subtypes, including enrichment of TNF-α/NF-κB, interferon response, and glycolysis-related pathways. **(B)** Heatmap showing scaled expression of differentially expressed genes across the same cell populations. Rows correspond to cell types, and columns represent genes associated with major enriched pathways. Patterns highlight cell type–specific expression programs, with fibroblasts enriched for extracellular matrix and TGF-β signaling genes, and malignant osteoblast subtypes showing heterogeneous activation of inflammatory and metabolic programs.

**Supplementary Figure 9.**
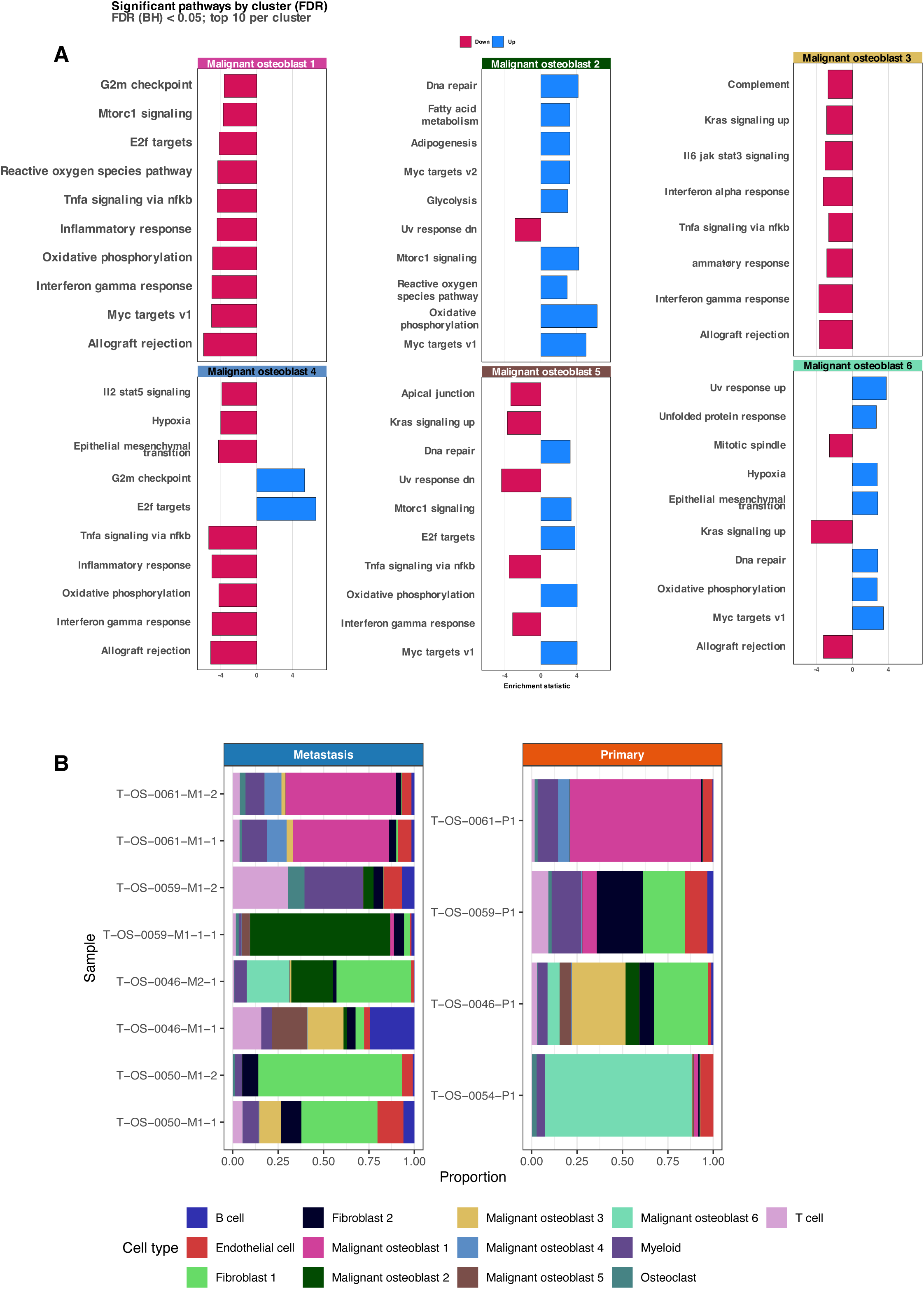
Functional pathway enrichment and cellular composition differences between malignant osteoblast subtypes and disease stages. **(A)** Gene set enrichment analysis using Wilcoxon-Mann-Whitney tests (wmw_gsea) for the six malignant osteoblast subtypes. Bars represent enrichment statistics for upregulated (red) and downregulated (blue) Hallmark pathways within each subtype (FDR < 0.05; top 10 per cluster). Distinct molecular programs are evident, including inflammatory and interferon signaling in malignant osteoblast 1, DNA repair and fatty acid metabolism in malignant osteoblast 2, and epithelial–mesenchymal transition and mitotic spindle pathways in malignant osteoblast 6, highlighting functional heterogeneity among tumor cell populations. **(B)** Stacked bar plots showing the proportional representation of annotated single-cell populations within metastatic (left) and primary (right) tumors from matched individuals. Malignant osteoblast subtypes dominated both stages, though primary tumors displayed higher prevalence of certain subtypes (e.g., malignant osteoblast 3), while metastatic samples exhibited increased heterogeneity and greater contributions from stromal and immune populations.

**Supplementary Figure 10.**
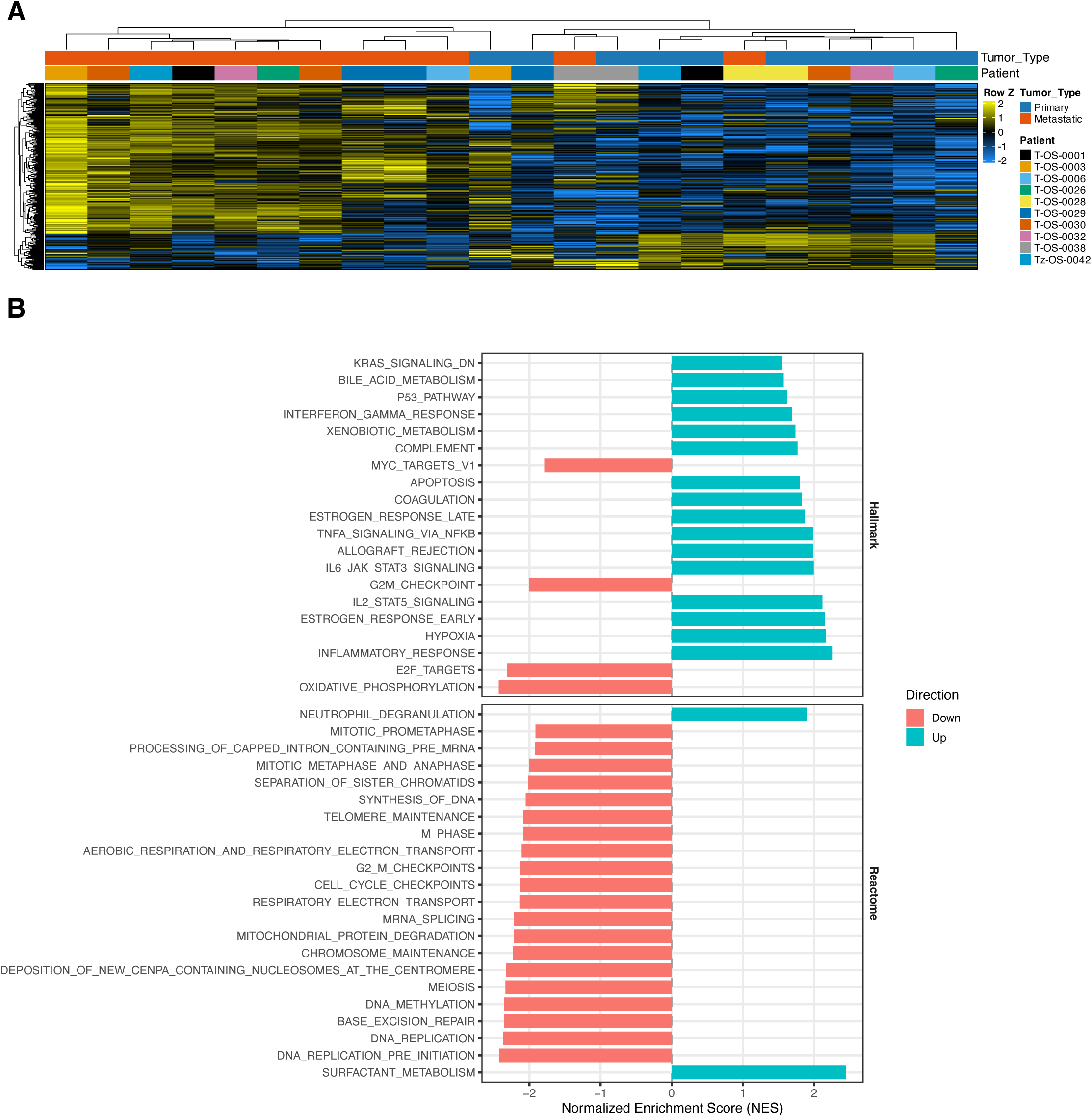
Differential expression and pathway enrichment from bulk RNA-seq analysis of primary and metastatic osteosarcoma. **(A)** Hierarchical clustering heatmap of differentially expressed genes (DEGs) between primary and metastatic tumors from bulk RNA-seq data. Each column represents an individual sample and each row a gene, with values Z-score–scaled by row. Clustering reveals patient-specific expression patterns alongside transcriptional differences between primary and metastatic tumors. **(B)** Pathway enrichment analysis of bulk DEGs using Hallmark (top) and Reactome (bottom) gene sets. Bars represent normalized enrichment scores (NES) for pathways upregulated (teal) or downregulated (pink) in metastases relative to primaries. Metastatic tumors show upregulation of interferon, inflammatory, and TNF-α/NF-κB signaling pathways, and downregulation of oxidative phosphorylation, cell-cycle, and DNA-replication pathways.

**Supplementary Figure 11.**
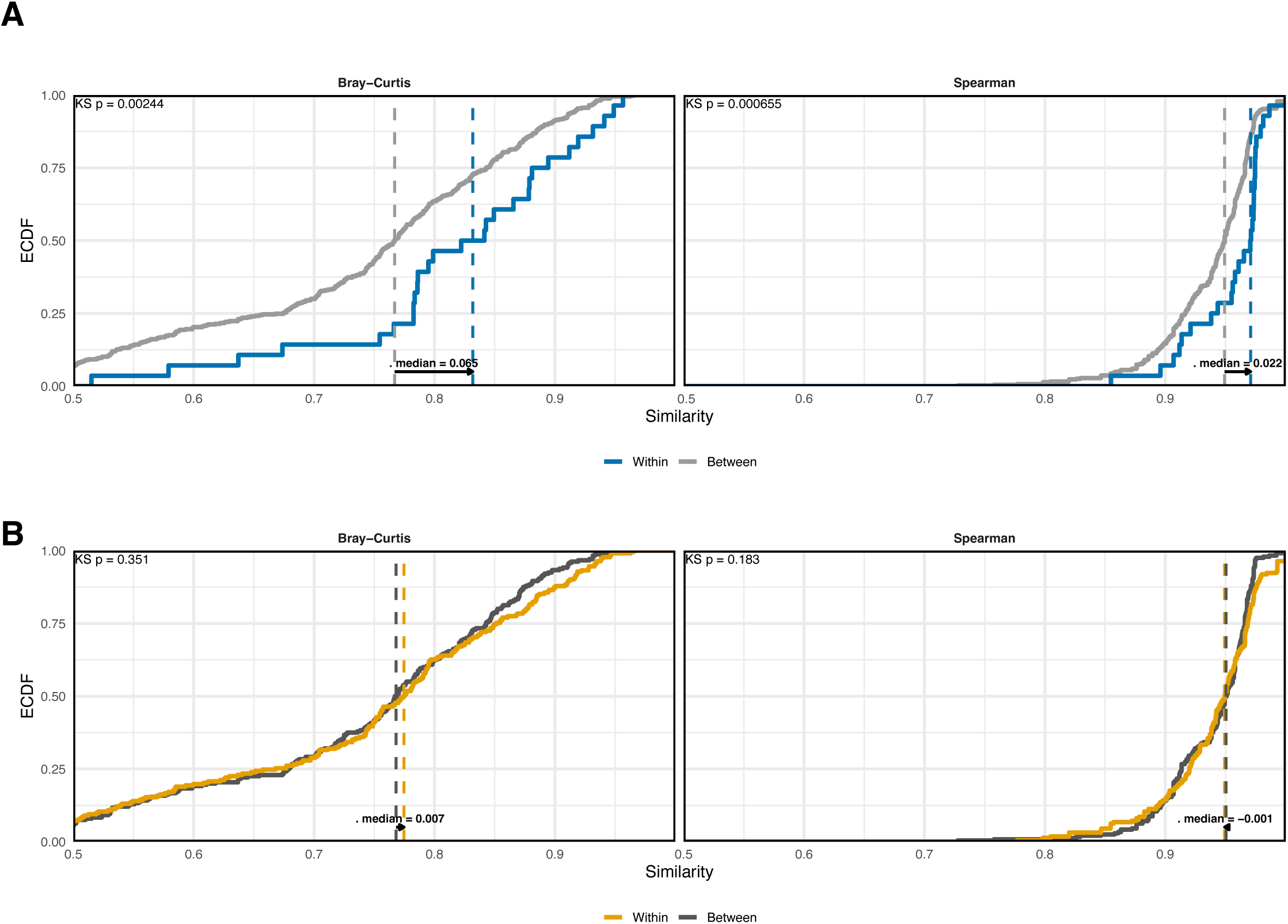
Bulk RNA-seq within- versus between-group similarity analysis. (A) Empirical cumulative distribution functions (ECDFs) comparing within- and between-patient similarity in bulk RNA-seq gene expression using Bray–Curtis (left) and Spearman correlation (right). Within-patient comparisons showed significantly higher similarity across both metrics (Bray–Curtis p=0.00244; Spearman p=0.000655), consistent with patient-specific expression profiles. (B) ECDFs comparing within- and between-stage (primary vs. metastatic) similarity using the same metrics. No significant stage-dependent differences were observed (Bray–Curtis p=0.351; Spearman p=0.183), indicating that transcriptional profiles are more strongly shaped by patient identity than disease stage.

**Tables**

n/a

